# De novo assembly and characterization of a highly degenerated ZW sex chromosome in the fish Megaleporinus macrocephalus

**DOI:** 10.1101/2024.03.07.583869

**Authors:** Carolina Heloisa de Souza Borges, Ricardo Utsunomia, Alessandro Varani, Marcela Uliano-Silva, Lieschen Valeria G. Lira, Arno J. Butzge, John F. Gomez Agudelo, Shisley Manso, Milena V. Freitas, Raquel B. Ariede, Vito A. Mastrochirico-Filho, Carolina Penaloza, Agustín Barria, Fábio Porto-Foresti, Fausto Foresti, Ricardo Hattori, Yann Guiguen, Ross D. Houston, Diogo Teruo Hashimoto

## Abstract

**Background:** *Megaleporinus macrocephalus* (piauçu) is a Neotropical fish within Characoidei that presents a well-established heteromorphic ZZ/ZW sex-determination system and thus, constitutes a good model for studying W and Z chromosomes in fishes. We used PacBio reads and Hi-C to assemble a chromosome-level reference genome for *M. macrocephalus*. We generated family segregation information to construct a genetic map, pool-seq of males and females to characterize its sex system, and RNA-seq to highlight candidate genes of *M. macrocephalus* sex determination.

**Results:** *M. macrocephalus* reference genome is 1,282,030,339 bp in length and has a contig and scaffold N50 of 5.0 Mb and 45.03 Mb, respectively. Based on patterns of recombination suppression, coverage, F_st,_ and sex-specific SNPs, three major regions were distinguished in the sex chromosome: W-specific (highly differentiated), Z-specific (in degeneration), and PAR. The sex chromosome gene repertoire was composed of genes from the TGF-β family *(amhr2*, *bmp7*) and Wnt/β-catenin pathway (*wnt4*, *wnt7a*), and some of them were differentially expressed.

**Conclusions:** The chromosome-level genome of piauçu exhibits high quality, establishing a valuable resource for advancing research within the group. Our discoveries offer insights into the evolutionary dynamics of Z and W sex chromosomes in fish, emphasizing ongoing degenerative processes and indicating complex interactions between Z and W sequences in specific genomic regions. Notably, *amhr2* and *bmp7* are potential candidate genes for sex determination in *M. macrocephalus*.

## 1. Background

The sex chromosomes constitute one of the most complex regions of the genome to sequence and assemble. The main challenges are associated with its haploid nature (high sequence divergence) and high repeat content. For instance, the Y chromosome in humans has been notoriously challenging to sequence and assemble due to its complex repeat structure [1], resulting in more than 50% of the chromosome missing from the reference assemblies [2]. Recently, the Telomere-to-Telomere (T2T) consortium presented a complete and gapless assembly of the human Y chromosome [3]. The new reference revealed, at single-base resolution, the previously uncharacterized 30 Mb of sequence within the long arm heterochromatic region. It also included newly assembled pseudoautosomal regions (PAR) and provided a full annotation of the gene, repeat, and organizational structure of the human Y chromosome [3]. Furthermore, the challenges faced during the assembly construction led to the development of novel automated methods for diploid genome assembly [4].

Unlike mammals and birds, teleost fish exhibit a vast range of sex-determination systems and mechanisms. Despite this, sex chromosomes are often overlooked in sequencing designs and when included, homogametic sexes (XX females or ZZ males) are generally favored. Currently, with the advance in sequencing technologies and the development of new bioinformatic tools, several Y chromosome assemblies have been reported in fish, revealing their remarkable diversity, *e.g.*, zig-zag eel [5], threespine stickleback [6], Atlantic herring [7] and the spotted knifejaw neo-Y [8].

A decade ago, the assembly of the first W chromosome in fish was conducted for the tongue-sole *Cynoglossus semilaevis* [9], offering a comprehensive understanding of its structure and evolution. To date, it stands as the only well-characterized W chromosome in fish. Despite being relatively young (approximately 30 million years old), the tongue-sole W chromosome exhibits a high rate of gene loss (70%), a small pseudoautosomal region (640 kb), suppression of recombination spread over most of the chromosome, and a high content of transposable elements (TE), which explains its larger size compared to the Z chromosome.

The genus *Leporinus* and *Megaleporinus*, within Anostomidae, comprise small to medium-sized fish, with straight body form and that present round stripes that can vary in number and size. *Megaleporinus* comprehends former *Leporinus* species that were placed in different genera due to the presence of the ZZ/ZW sex chromosome system [10]. *Leporinus* has homomorphic sex chromosomes, whereas *Megaleporinus* sex chromosomes are heteromorphic and have existed for at least 12 million years [10].

The piauçu *Megaleporinus macrocephalus* is the only aquaculture species of Brazil (South America) with a well-established ZZ/ZW heteromorphic sex chromosome system [11]. The W chromosome of the species is the largest of the karyotype and presents a huge C-positive heterochromatic block occupying the entire long arms, full of repetitive DNA [12]. The Z chromosome is a medium-sized metacentric with only portions of heterochromatin at the end of the long arms [11].

The study of W chromosome evolution in vertebrates continues to be constrained by the scarcity of W assemblies; therefore, it is of paramount importance to sequence additional W models. *Megaleporinus macrocephalus* is an ideal model for studying W chromosome structure and evolution in fish, as it belongs to a rare group with conserved ZW chromosomes. In this study, we aimed: 1) to assemble a chromosome-level genome of *Megaleporinus macrocephalus*, including the sex chromosome; 2) to create a linkage mapping for the species to assess patterns of recombination; 3) to carry out resequencing of male and female individuals to identify sex-linked regions in the genome; and 4) to conduct RNA-seq experiment to identify candidate genes for sex determination.

## 2. Results

### 2.1 Chromosome-level genome assembly

#### 2.1.1 Genome Assembly

We generated 88.8 Gb of Pacific Biosciences (PacBio) continuous long reads (CLR), 85 Gb of MGISEQ short reads, and 105 Gb of Hi-C data. Genome coverage based on final assembly size was 69.4x, 66.4x, and 82x, respectively. The unique molecular yield of PacBio reads was 56 Gb and the subread N50 length was 32 kb. Regarding the short reads, after the removal of poor-quality sequences, we kept 82 Gb of clean data. This dataset was used to generate *k*-mer spectrum plots to estimate the overall characteristics of the genome. All *k*-mer plots were similar and showed a profile correspondent with a low heterozygosity rate (**Figure 1A**). The estimated genome size (21-mer) was 1.04 Gb with a heterozygosity of 0.48% and 18% of repeat content.

**Figure 1.**
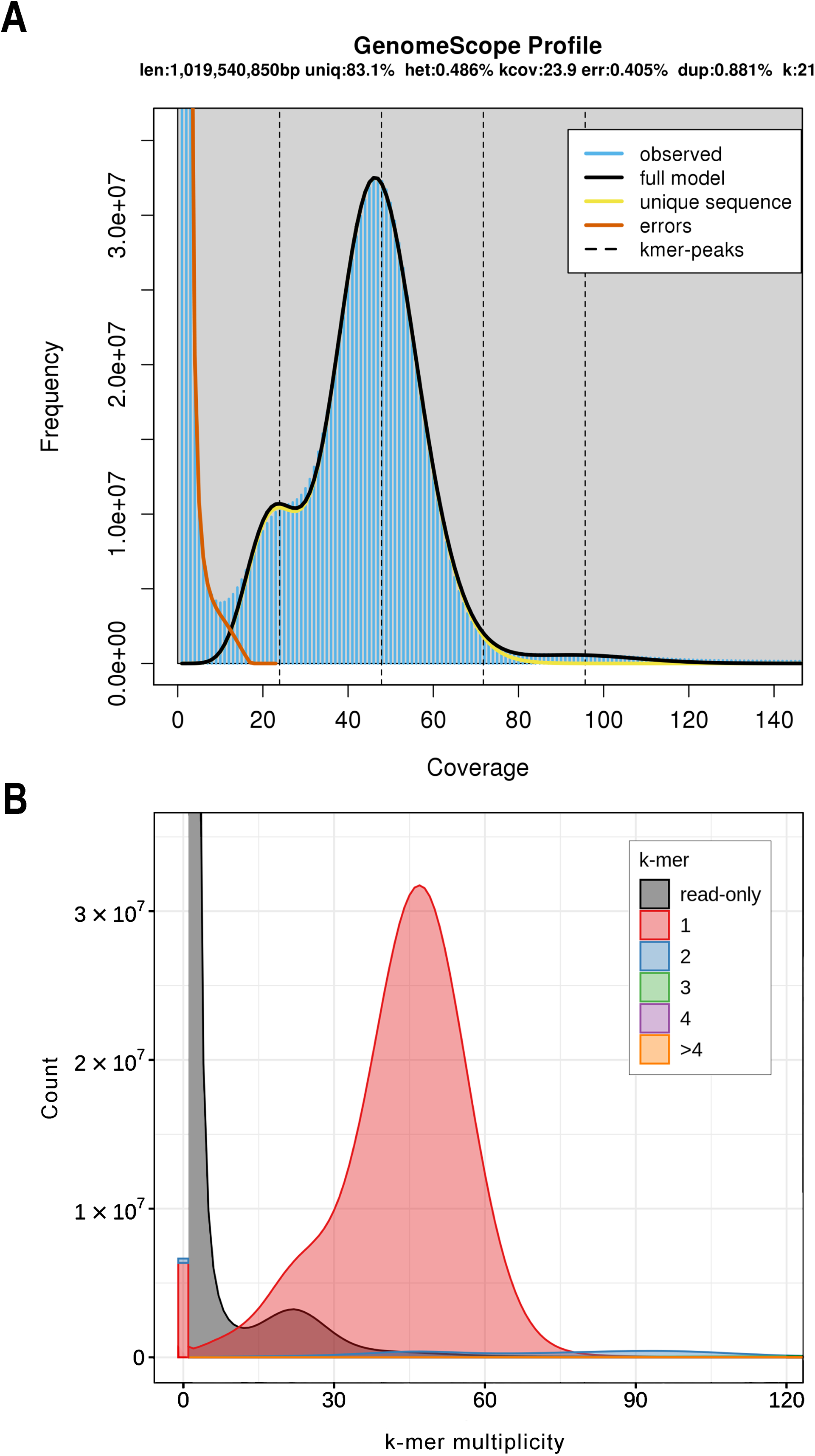
*K*-mer profile of MGISEQ short reads (A). A *k*-mer analysis of the *Megaleporinus macrocephalus* genome bases against its sequenced MGISEQ reads (B).

We used Falcon/ Falcon-Unzip [13][14], Flye [15][16], wtdbg2 [17][18], and Canu [19][20] to assemble the PacBio long reads. Falcon/ Falcon-Unzip [13][14] assembly presented the best contiguity metrics (2,770 primary contigs, 33 contigs > 5 Mb, with N50 of 1.53 Mb) and was chosen for further analysis. After gap filling, the initial contigs were clustered in 1,227 scaffolds with N50 of 5.0 Mb. The scaffolds were ordered and oriented into 27 chromosomes, which is consistent with the haploid chromosome number of the species [21]; and 73 unplaced scaffolds (< 250 kb). The 27 chromosomes comprised 99.56 % of the complete genome assembly. The final *M. macrocephalus* reference genome contains 27 chromosomes and 73 unplaced scaffolds. It has a contig and scaffold N50 of 5.0 Mb and 45.03 Mb, respectively, and an assembled genome size of 1.28 Gb (**Table 1**).

**Table 1.** Statistics for genome assembly of *Megaleporinus macrocephalus*.

| Characteristic | Value |
| --- | --- |
| No. scaffolds | 101 |
| No. contigs | 1,353 |
| Main genome scaffold sequence total (bp) | 1,282,030,339 |
| Main genome contig sequence total (bp) | 1,280,781,659 |
| Scaffold N50 (bp) | 45,034,219 |
| Contig N50 (bp) | 5,013,076 |
| Max. scaffold length (bp) | 73,843,892 |
| Max. contig length (bp) | 25,940,738 |
| % main genome in scaffolds > 50 kb | 99.9% |
| BUSCO complete | 96.2% |
| BUSCO complete and single copy | 95.1% |
| BUSCO complete and duplicated | 1.1% |
| BUSCO fragmented | 0.6% |
| BUSCO missing | 3.2% |
| Consensus quality value (QV) | 37.53 |
| Mercury completeness | 93.05% |

We used the highly accurate short reads to plot Merqury [22] [23] evaluation against the genome *k*-mers. **Figure 1B**, shows that (i) the distribution of the *k*-mers in the assembly is consistent with the short read profile (**Figure 1A**), (ii) two peaks are demonstrating that 1-copy (heterozygous) and 2-copy (homozygous) *k*-mers were found once in the assembly, as expected for a pseudo-haplotype genome [23], (iii) most of the assembly *k*-mers (in red) are unique, indicating that the assembly has a low content of artificial duplications (*i.e.*, *k*-mers found twice, in blue) (iv) there are missing *k*-mers in the assembly (black peak), which is compatible with haploid genomes, (v) the 1-copy *k*-mer peak (red) is greater than its missing sequences (black), this suggests that Falcon-Unzip [14] erroneously included sequences from both haplotypes into the primary pseudo-haplotype [23]. Also, this possibly led to an assembled genome size greater than the estimated (1.04 Gb). The accuracy of the base calls (QV), which is calculated using the *k*-mers found only in the assembly (bar at the beginning of **Figure 1B**), was 37. 53 (**Table 1**) and represents a base accuracy > 99.9% (*e.g*., QV = 30 means 99.9% accuracy). The completeness score shows that 93.05% of *k-*mers in the MGISEQ reads are present in the assembly, which is a good recovery of *k*-mers for a species with 0.5% heterozygosity.

Pearson’s correlation between the autosomes assembled size with its actual karyotypic size (**Supplementary Table 1**) was 99%, demonstrating the high quality of the assembled *M. macrocephalus* genome.

#### 2.1.2 Sex chromosome

Chromosome 13 was recognized as the sex chromosome based on the following evidence:

- In the Hi-C contact map, we observed a lower coverage in the upper segment of this chromosome compared to its terminal segment and other chromosomes. We assumed that this segment corresponds to the W-specific region (hemizygous), and the terminal segment corresponds to the pseudo-autosomal region of Z and W (**Figure 2**).

**Figure 2.**
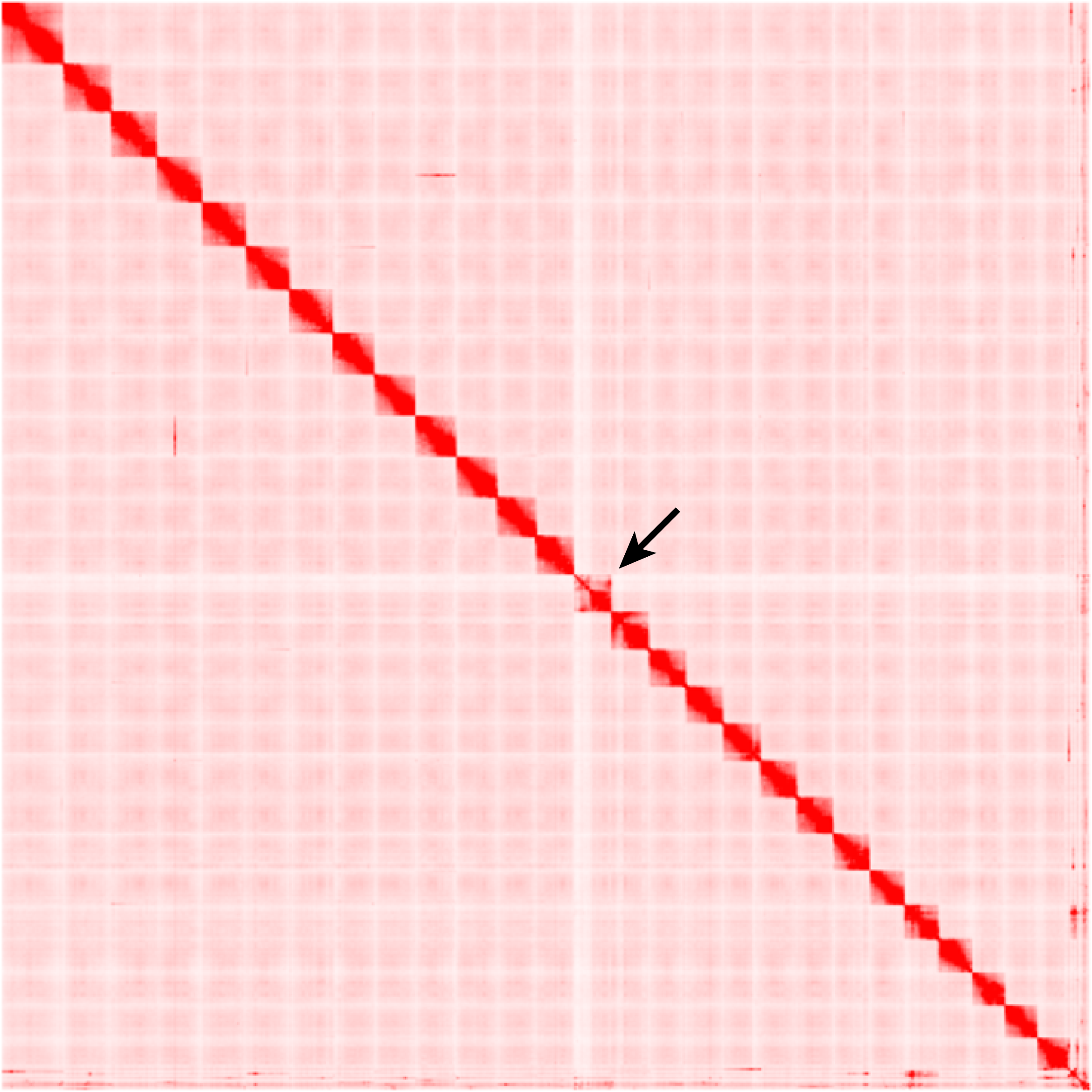
Hi-C contact map highlighting the sex chromosome (arrow) of *Megaleporinus macrocephalus*.

- In the linkage map, linkage group (LG) 24 (= chromosome 13) exhibited suppression of recombination, with varying intensities in the female and male maps (**Figure 4)**.

- The comprehensive examination of SNP distribution through resequencing analysis unveiled a robust sex-linked signal in females and elevated fixation index (*F*_ST_) values within chromosome 13 **(see Figure 5).**

#### 2.1.3 Repeat Annotation

Using the *de novo* prediction model, 2,544 new families of repeats were found in the genome. The repeat content found in *M. macrocephalus* accounted for 46.71% of the genome (598 Mb). Among the repeats, transposable elements were the most common representing 37.49% of the genome. DNA transposons were the most abundant TE (11.82%), following 3.02% of long terminal repeats (LTR), 3.42% of long interspersed nuclear elements (LINE), and 0.33% of short interspersed nuclear elements (SINE) (**Supplementary Table 2**). A great percentage (18.89%) of the interspersed repeats remained unclassified. Despite using the satellitome of the species [12] to identify the sat DNAs of the genome, these repeats accounted for only 4.40% of the genome (**Supplementary Table 2**).

**Supplementary Figure 1** shows older TE copies located on the right side of the graph and rather recent ones, that do not diverge much from the consensus TE sequence, on the left side. Most of the interspersed repeat content found in the *M. macrocephalus* genome is recent (*K*-values < 25). Also, it is possible to observe two bursts of transposition dominated by DNA transposon.

The repeat content found in the sex chromosome was slightly higher than in the autosomes (4.24%). Total interspersed repeats and satellites were the classes that presented the major difference, 2.37% and 2.24%, respectively (**Table 2**).

**Table 2.** Comparison between the repeat content in the sex chromosome and the autosomes of the *Megaleporinus macrocephalus* genome.

|  | % sex chromosome | % autosomes |
| --- | --- | --- |
| **Repeat content** | 50.95 | 46.71 |
| **Retroelements** | 7.86 | 6.78 |
| **DNA transposons** | 12.07 | 11.82 |
| **Unclassified** | 19.92 | 18.89 |
| **Total interspersed repeats** | 39.86 | 37.49 |
| **Small RNA** | 0.02 | 0.03 |
| **Satellites** | 6.64 | 4.40 |
| **Simple repeats** | 3.72 | 4.06 |
| **Low complexity** | 0.38 | 0.40 |

#### 2.1.4 Gene Prediction and Annotation

For *ab initio* gene prediction, BRAKER1 [24] [25] used 28.26 Gb of RNA-seq data as extrinsic evidence to predict 60,482 genes. For homology-based gene prediction, BRAKER2 [26] [25] generated 57,574 hints and predicted genes. TSEBRA [27] [28] combined BRAKER runs and selected 44,054 best gene predictions. Of these, 66.94% (29,490) were annotated in the Actinopterygii database of Eggnog or UniProtKB [29]; and 33.06% (13,525) were not annotated. We kept the annotated (29,490) and the non-annotated predicted genes with more than 150 amino acids (1,039) for the final dataset, summarizing 30,501 protein-coding predicted genes (**Supplementary Table 3**). The final dataset had 94.1% complete Benchmarking Universal Single-Copy Orthologs (BUSCO), 89% complete and single-copy, 5.1% duplicated, 2.2% fragmented, and 3.7% missing BUSCO. For the functional annotation, we performed blast searches against the Actinopterygii database of UniProtKB [29]. Of all the predicted genes, only 3.34% (1,018) were not annotated.

The most representative gene ontology (GO) terms (> 15% of genes) according to the three-domain can be seen in **Supplementary Figure 2**.

### 2.2 Linkage map

A total amount of 1,307,500,332 raw reads were sequenced by Double Digest Restriction Site-Associated DNA Sequencing (ddRADseq), resulting in approximately 200 Gb of data (approximately 28 Gb per library). After filtering (removal of low-quality sequences and reads with missing or ambiguous barcodes), an average of 11% of the reads were removed from each library, *i.e.*, 89% of the reads were retained for analysis. Furthermore, 24 individuals were excluded due to the low number of reads (< 1 million). The average number of reads per sample was 4.3 million. Raw sequencing data and filtered reads for each library are shown in **Supplementary Table 4**.

After mapping the ddRAD reads to the chromosome-level genome, SNP calling analysis resulted in 41,033 SNPs from 85,167 loci from 281 individuals. In Plink [30] [31], after applying the mind and geno filters, 56 individuals and 8,971 SNPs were excluded. At last, the maf filter excluded 3,733 SNPs. Thus, 225 individuals and 28,329 SNPs passed all quality controls (total genotyping rate of 0.96) and were used for the linkage mapping.

We performed a pedigree test and individuals with > 10% of Mendelian errors were removed. After calling possible missing or erroneous parental genotypes in the *ParentalCall* module, a total of 9,997 SNPs were joined to linkage groups (LGs). We computed several Logarithm of Odds (LOD) scores between markers and selected the best marker distribution according to the karyotype characteristics of the species. Although the *M. macrocephalus* haploid chromosome number is 27, the best distribution of markers was achieved using 28 LGs (with LOD 12), probably because of the specific region of the Z chromosome that constitutes a separate linkage group (**Supplementary Figure 3)**. The remaining markers were assigned to the existing LGs using LOD 10, which recovered 1,234 markers. 18,098 markers were discarded because no association with the linkage map was detected. In each LG, the orders of the markers with the best likelihood were combined to produce the final linkage map. A total of 11,231 SNPs were assigned to 28 LGs. We constructed a male, a female, and a sex-averaged map (average position between male and female map) (**Figure 3**).

**Figure 3.**
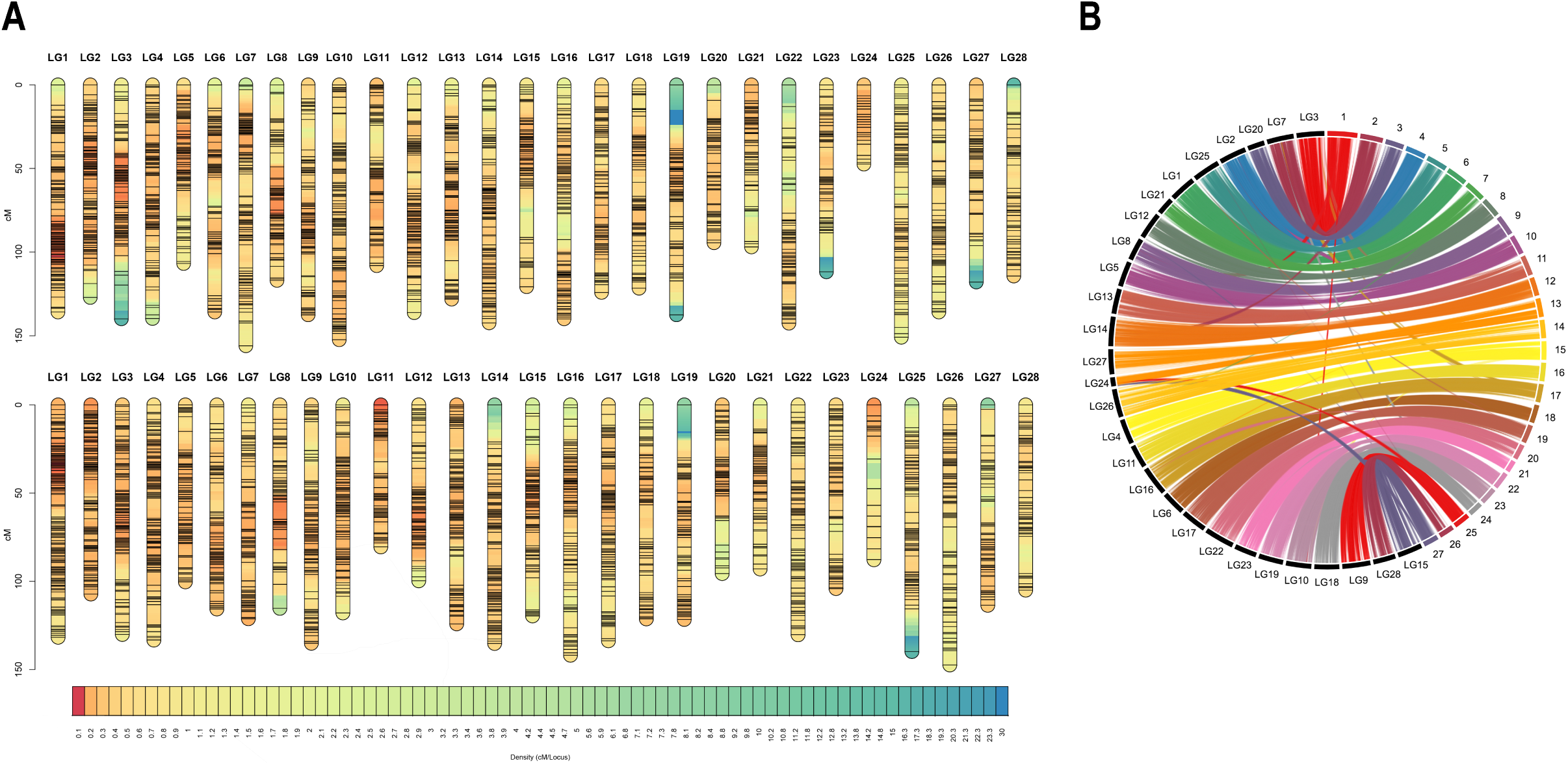
Male (A) and female (B) linkage maps of *Megaleporinus macrocephalus* showing 28 linkage groups and 11,231 SNPs. The density of markers is represented by a range of different colours that vary from blue (low density regions) to red (high density regions).

The number of SNPs in the LGs varied from 710 (LG1) to 203 (LG28). In the sex-averaged map, LGs length ranged from 143.08 (LG22) to 43.25 (LG24) centimorgans (cM), with an average of 3320.36 cM and an average distance between markers of 0.29 cM (SD = 0.12). The highest and lowest marker densities were found on LG1 and LG22, with an average of 0.18 and 0.61 cM, respectively (**Supplementary Table 5**).

Concerning sex-specific differences, the average distance between markers in male and female maps were 0.31 and 0.29, respectively. Therefore, the male map (3,518.24 cM) was longer than the female (3,301.97 cM). The male:female genetic length ratio over the entire genome was 1.07. The ratios varied from 0.54 (LG24) to 1.37 (LG12). The highest density of recombination was detected at the proximal region of the centromeres (considering that this species has metacentric/submetacentric chromosomes), although some exceptions occurred in the terminal region of the LG11 (**Figure 3**).

#### 2.2.1 Recombination suppression within LG24

In LG24, recombination was distributed differently between the sexes (heterochiasmy). In the female map, LG24 was almost double the size (87.81 cM) of the same LG in the male map (47.37 cM). The comparative synteny analysis revealed a correspondence of LG24 with the sex chromosome 13 (**Figure 3**), particularly in regions < 20 Mb and > 40 Mb. Also, zero recombination clusters (suppression of recombination) were observed in this LG for both sexes (**Figure 4**), *i.e.*, blocks of markers that vary in the physical distance (bp) but do not vary in the genetic distance (cM). Besides, in the genomic synteny between LG24 and the sex chromosome 13 (**Figure 3**), the same chromosome was also attributed to LG27, probably in the PAR. This indicates that the best LOD value resulted in 28 LGs (n = 27 chromosomes) because both LG24 and LG27 corresponded to the same chromosome (different regions of the sex chromosome).

**Figure 4.**
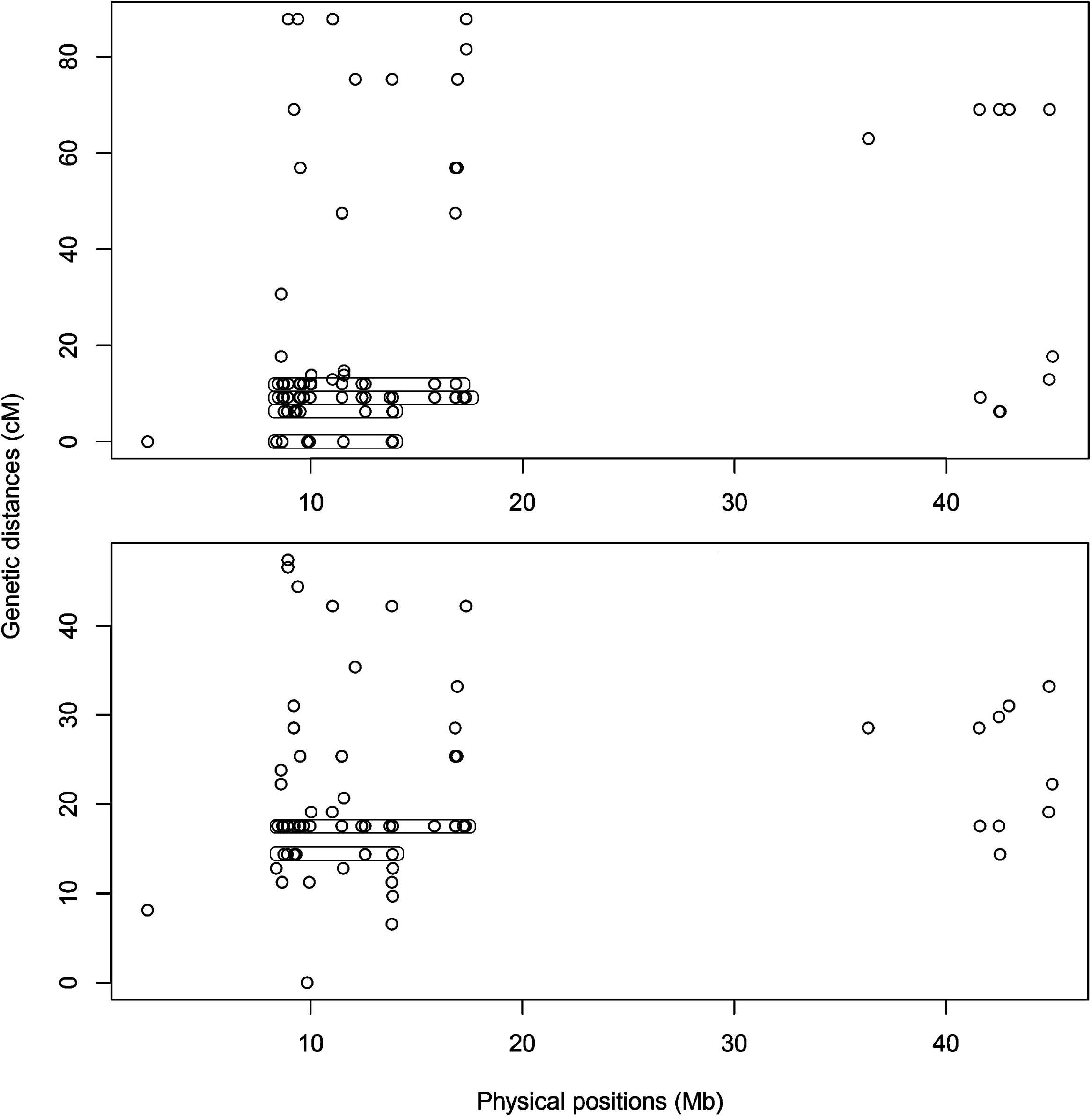
Marey maps of LG24. Female map at the top, male at the bottom. Recombination varies in the physical map but not in the genetic map forming vertical structures of clusters (zero recombination clusters), highlighted by circles.

#### 2.2.2 Discordance between physical and genetic mapping

To integrate (reconcile) the genome assembly with the linkage map data, we ordered the genome scaffolds using the linkage map as a reference. Chromonomer [32] [33] tries to identify and remove markers that are out of order in the genetic map when considered against their local assembly order; and to identify scaffolds that have been incorrectly assembled according to the genetic map, and split those scaffolds. The ordering grouped 1,575 map markers in 352 scaffolds. The remaining SNP loci were not used for genome anchoring because they were not aligned to the piauçu scaffolds or were markers mapping to multiple regions or loci where the orientation could not be suitably assigned. These results allowed the construction of a chromonome that clustered 75% (1,221,855,406 bp) of the initial scaffold data into 27 pseudomolecules (chromosomes) totalizing 977 Mb of length. 320 Mb were not anchored in pseudomolecules (chromosomes). LG24 anchored a low number of scaffolds, resulting in a pseudomolecule with poor scaffolding and small size (∼ 4 Mb). This can be explained by the region of sex conflict between the Z and W chromosomes and, consequently, the suppression of recombination between them.

The dot plot synteny analysis demonstrated a high degree of concordance between the chromosomes scaffolded with Hi-C data (physical mapping) and the linkage groups of the genetic map (**Supplementary Figure 4**, illustrated by chromosomes 5, 8, and 20). Insertions and deletions were observed in all chromosomes (*e.g.*, chromosome 2 since 25% of the initial scaffold data was missing). Beyond that, structural differences between the linkage map and scaffolds were noted in some chromosomes, revealing relocations (chromosomes 1 and 21) and major inversions (chromosomes 3, 18, and 19).

### 2.3 Highly differentiated regions in ZW chromosome

Whole genome sequencing of male and female pools yielded respectively 266,697,484 and 231,722,384 paired-end clean reads in total. Subsequently, the reads were mapped to the female chromosome-level genome to characterize genomic regions enriched for sex-biased signals. *i.e*., sex coverage differences or sex-biased SNPs. The mapping rates of paired-end reads from the male pool and female pool were 98.74 % and 97.08 % respectively, and the average depth of the male pool and female pool were 25 and 24, respectively.

The analysis of SNP distribution revealed a strong sex-linked signal in females and high *F*_ST_ values in the sex chromosome (chr 13) (**Figure 5**). This profile illustrates a female heterogametic system (ZW/ZZ), as previously reported by cytogenetic data [11].

**Figure 5.**
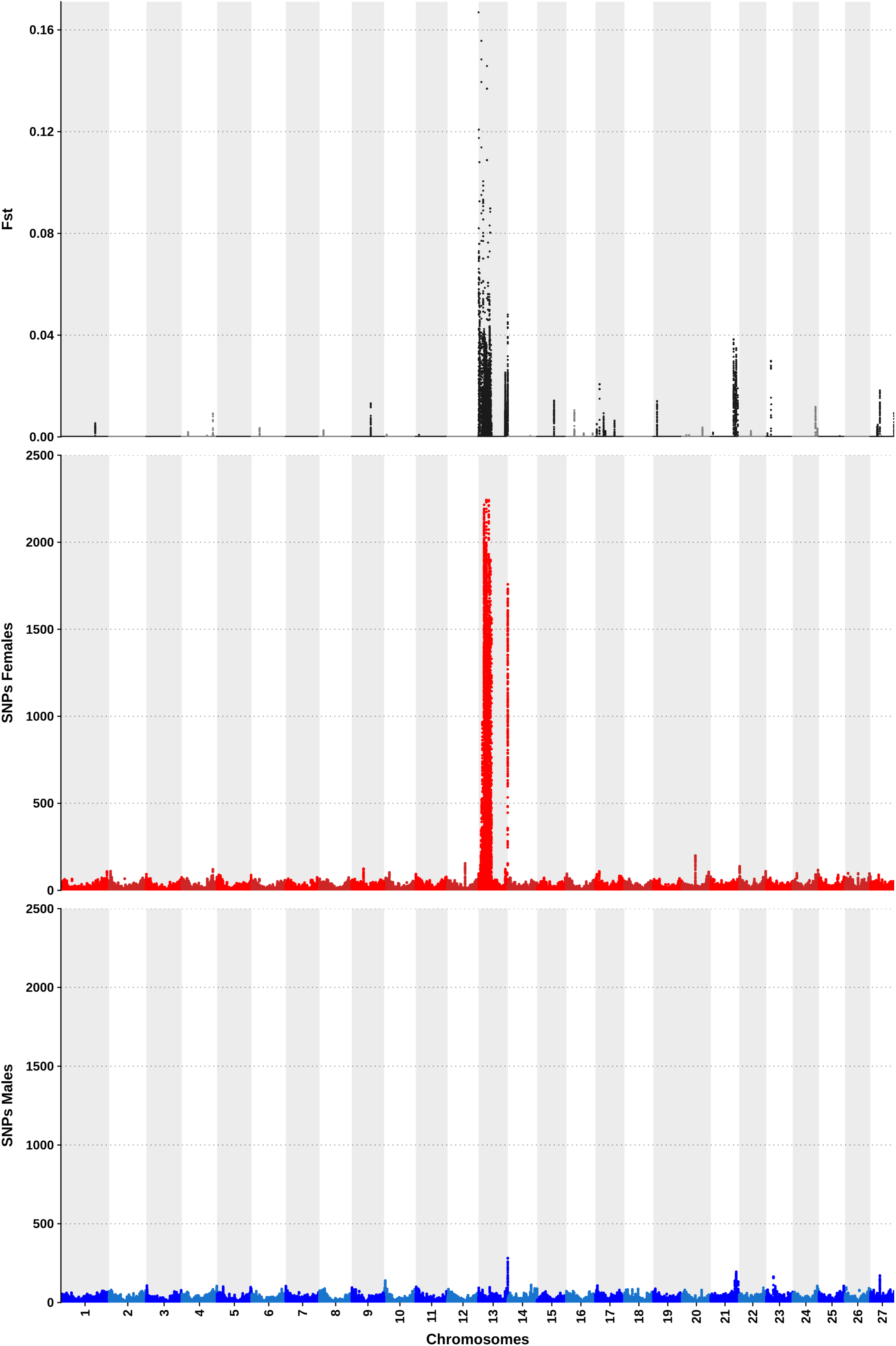
Plots of *F*_ST_, female and male-specific SNPs, respectively, accounted through windows of 50 kb along the 27 chromosomes of *Megaleporinus macrocephalus* genome.

#### 2.3.1 Distinct patterns in the sex chromosome

The sex chromosome, which is approximately 45 Mb in length, was divided into three regions according to the overall characteristics of read depth (coverage), the pattern of sex-specific SNPs, and *F*_ST_. The first region comprises the beginning of the chromosome, from 0 to ∼ 3 Mb and it is characterized by high coverage in females (2.3-fold the female average depth), and low coverage in males (0.6-fold the male average depth, **Figure 6**). The absence of coverage in males was detected in some areas (depth ratio ≍ 0). In addition, the major *F*_ST_ peak (*F*_ST_ = 0.17) was located within this region (**Figure 6**). The observed patterns strongly suggest the assembly of W-specific sequences in this region. However, Z-specific sequences are also present, albeit in much lower quantities (depth ratio > 1). This region was named a putative W-region (PWR), characterized by high differentiation, as confirmed by the absence of recombination in the linkage map (**Figure 4**).

**Figure 6.**
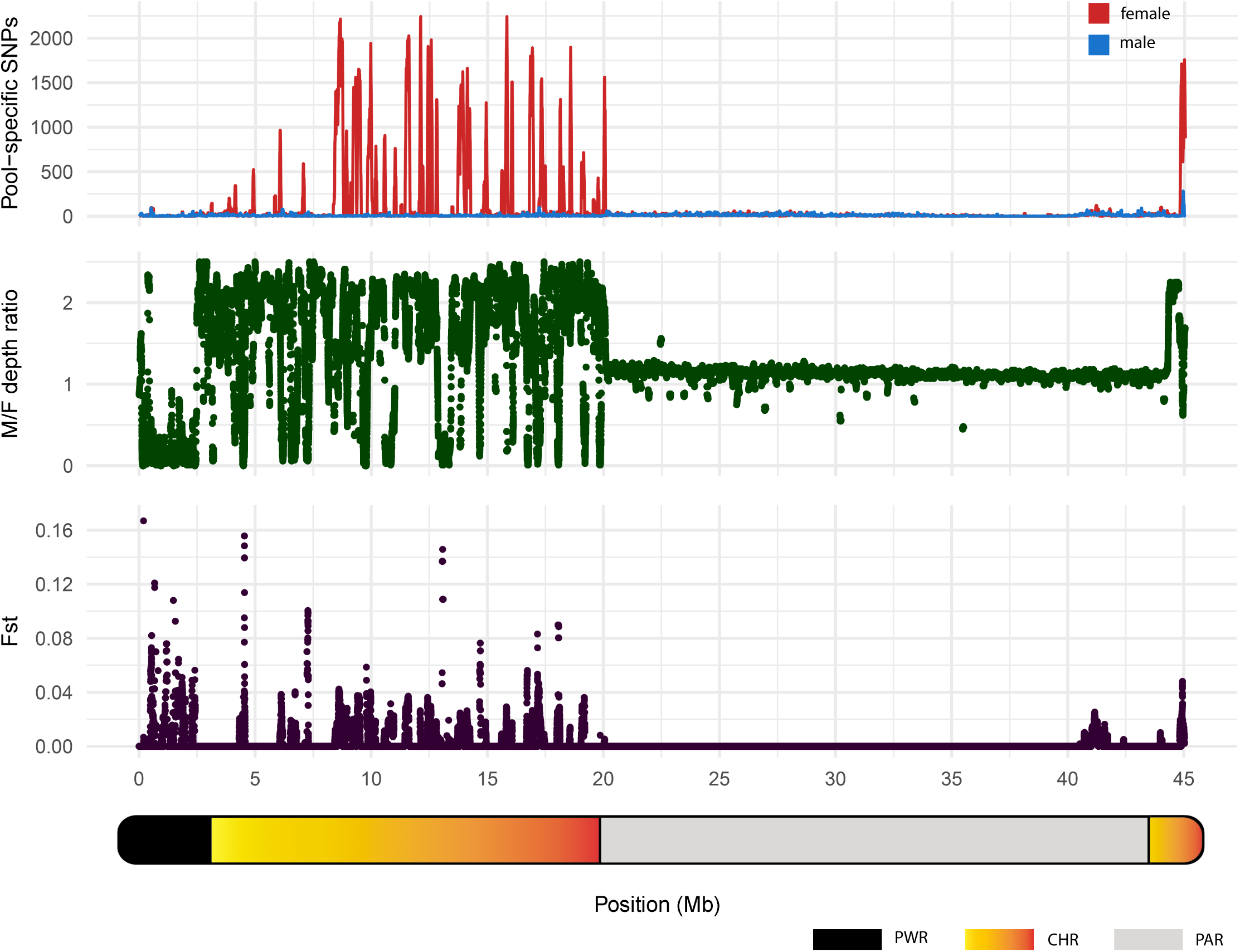
Pool-specific SNPs, M/F depth ratio (absolute depth males/ absolute depth of females) and *F*_ST_ along the sex chromosome of *Megaleporinus macrocephalus*. Depth ratio of = 1 indicates equal read coverage in males and females, while depth ratio > 1 and < 1 mean superior coverage in males ZZ (duplicated chromosome) and females ZW (single copy of each chromosome), respectively.

The second region encompasses ∼ 3 to 20 Mb, reaching the opposing terminal segment of the sex chromosome, which ranges from ∼ 44 to 45 Mb (**Figure 6**). Within this zone, two distinct patterns were recognized. The first, prevalent in most of the region, exhibited Z-specific characteristics, as males demonstrated at least double the coverage of females (*i.e.*, males have two copies of Z, while females have one). The other pattern was characterized by a high density of female-specific SNPs (with peaks summarizing more than 2,000 SNPs), representing allelic differences between Z and W sequences.

Furthermore, in the areas where W sequences were observed, males had no coverage, and females had 2.6-fold the average depth, resulting in a depth ratio ≍ 0 (**Figure 6**). This observation was supported by a higher recombination frequency in males compared to females within this region of LG24 (3 to 20 Mb and 44 to 45 Mb, as illustrated in **Figure 4**). This evidence indicates a certain degree of similarity between the Z and W sequences, allowing them to be scaffolded in the same region. Therefore, this locus was named ‘chimera’ (CHR), which is undergoing degeneration.

The region comprising ∼20 to 44 Mb was characterized by a lack of sex-specific SNPs (**Figure 6**). In this genomic locus was also seen an almost equal absolute depth between males and females (depth ratio ≍ 1, **Figure 6**). This illustrates homology between the male and female sequences in this zone and could indicate normal recombination rates as seen in pseudo-autosomal regions. Therefore, we named this region as PAR.

### 2.4 Differential expression between males and females

A total amount of 28.26 Gb of gonadal paired-end RNA-seq data was pseudo-aligned with 30,500 transcripts of *M. macrocephalus* (**Supplementary Table 6)**. Approximately 99.9% (30,460) of RNA-Seq transcripts were successfully pseudo-aligned, and, after low counts were filtered (≤ 1), 27,120 transcripts remained for the differential expression analysis. Principal component analysis (PCA) showed that 78% of the variance in the data was explained by Principal component 1 (PC1). Throughout PC1 the samples were clustered in two groups, ZZ males and ZW females, as expected (**Figure 7A**), despite being observed a minor intra-variation in the former. The heatmap of the Euclidean distance matrix demonstrated the same pattern (**Figure 7B**).

**Figure 7.**
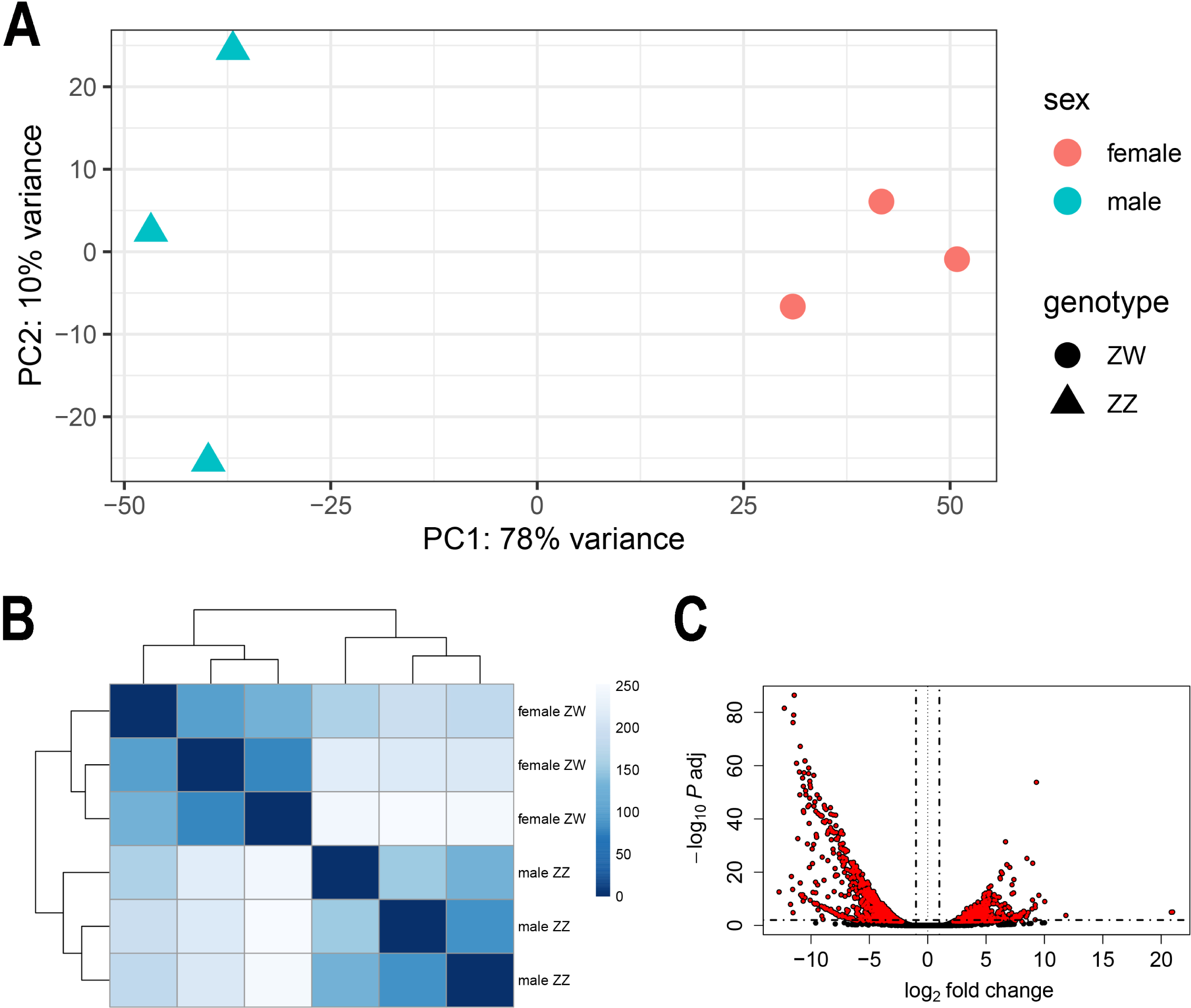
RNA-Seq samples clustered according to their transcript expression. PCA (A) and Euclidean distance matrix (B). Volcano plot showing down and up-regulated transcripts in red on the left and right, respectively. Not significantly expressed transcripts (*p*adj > 0.05) in black (C).

The analysis resulted in 2,557 differentially expressed transcripts (*p*adj ≤ 0.01); of which 42.66 % (1,091) were up-regulated in males, and 57.33% (1,466) up-regulated in females (**Table 3 and Figure 7C**). Most of the differentially expressed transcripts were components of the zona pellucida and up-regulated in females (**Supplementary Table 7**).

**Table 3.**
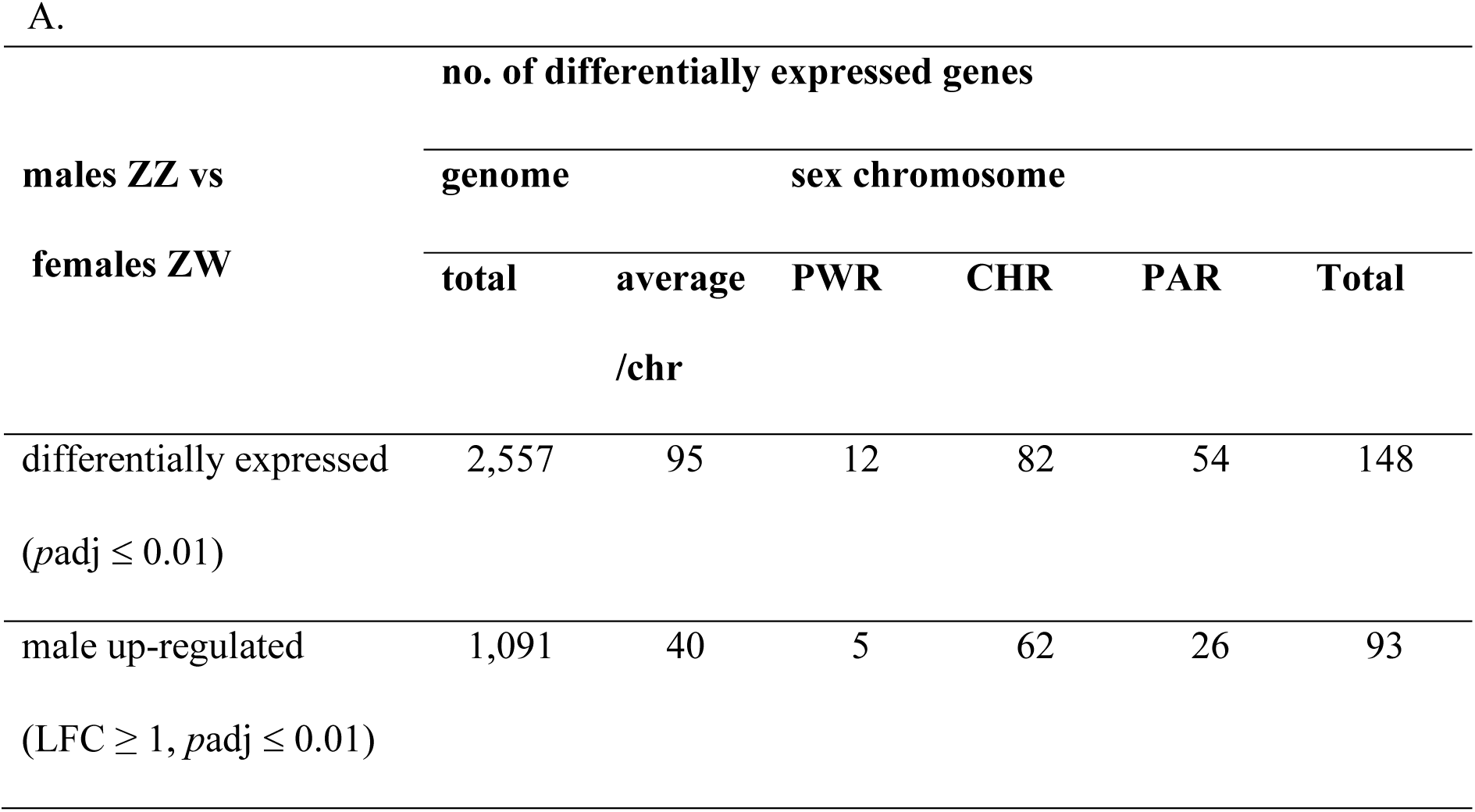
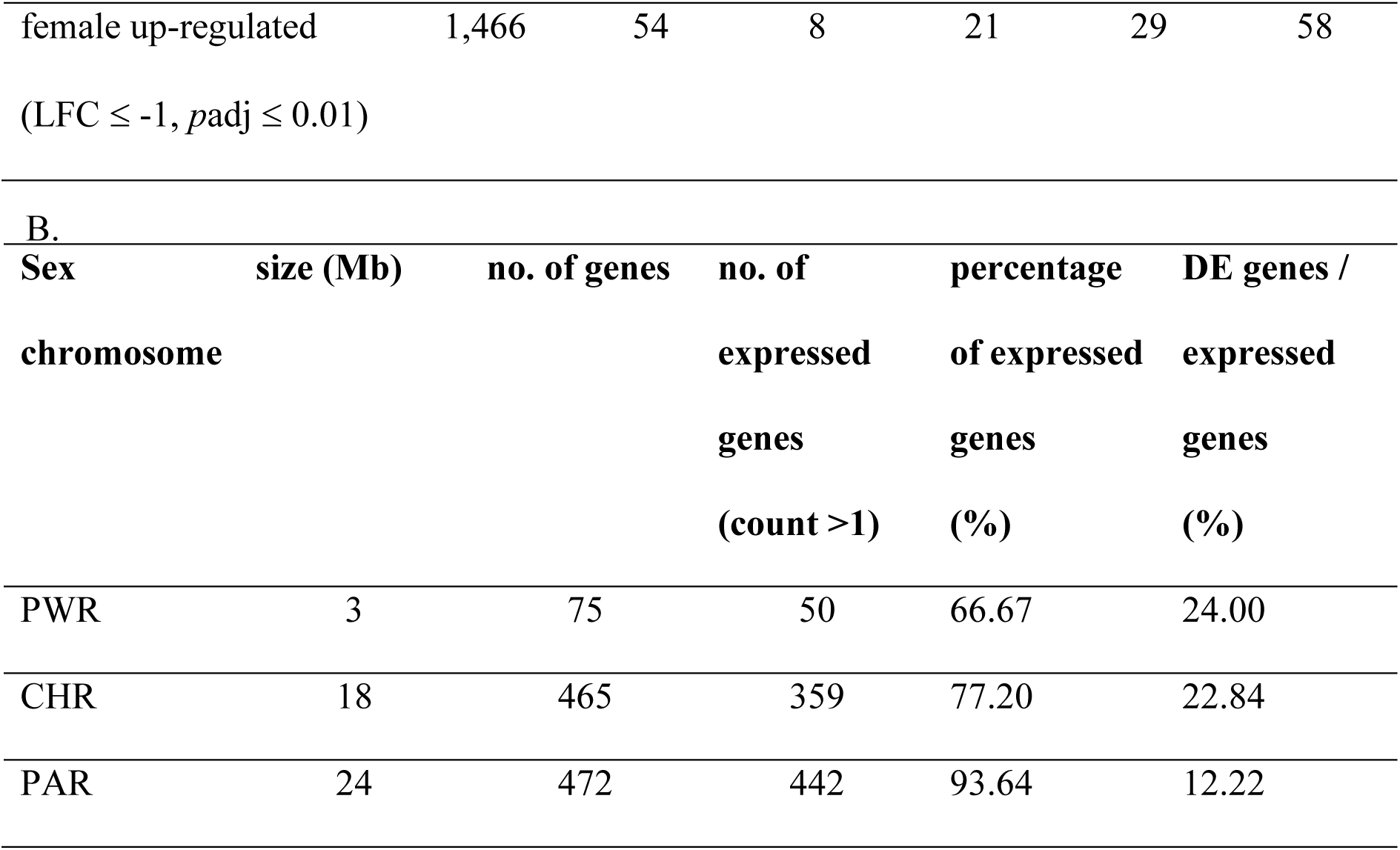
Summary of differentially expressed genes of males ZZ and females ZW (A) and within the sex chromosome (B) of *M. macrocephalus*.

In the sex chromosome, males were more up-regulated and presented more differentially expressed (DE) genes than the average per chromosome. Within it, the PWR was the richest in DE genes. Males were significantly more up-regulated in CHR than females (males are homozygous for Z and present great gene expression), which was also observed in PWR and PAR (**Table 3A).**

The PWR had the lowest percentage of expressed genes compared to the others, indicating the presence of genes that lost their functions. Despite it having only 3 Mb, 24% of its expressed genes were differentially expressed (**Table 3B**). In contrast, PAR, which has 24 Mb, presented only 12.22%. The lower expression in PWR and CHR compared to PAR indicates that genes in those regions may have suffered some degree of degeneration (**Table 3B**).

The sex chromosome presented several sex determination-related genes, as seen in **Table 4**.

**Table 4.**
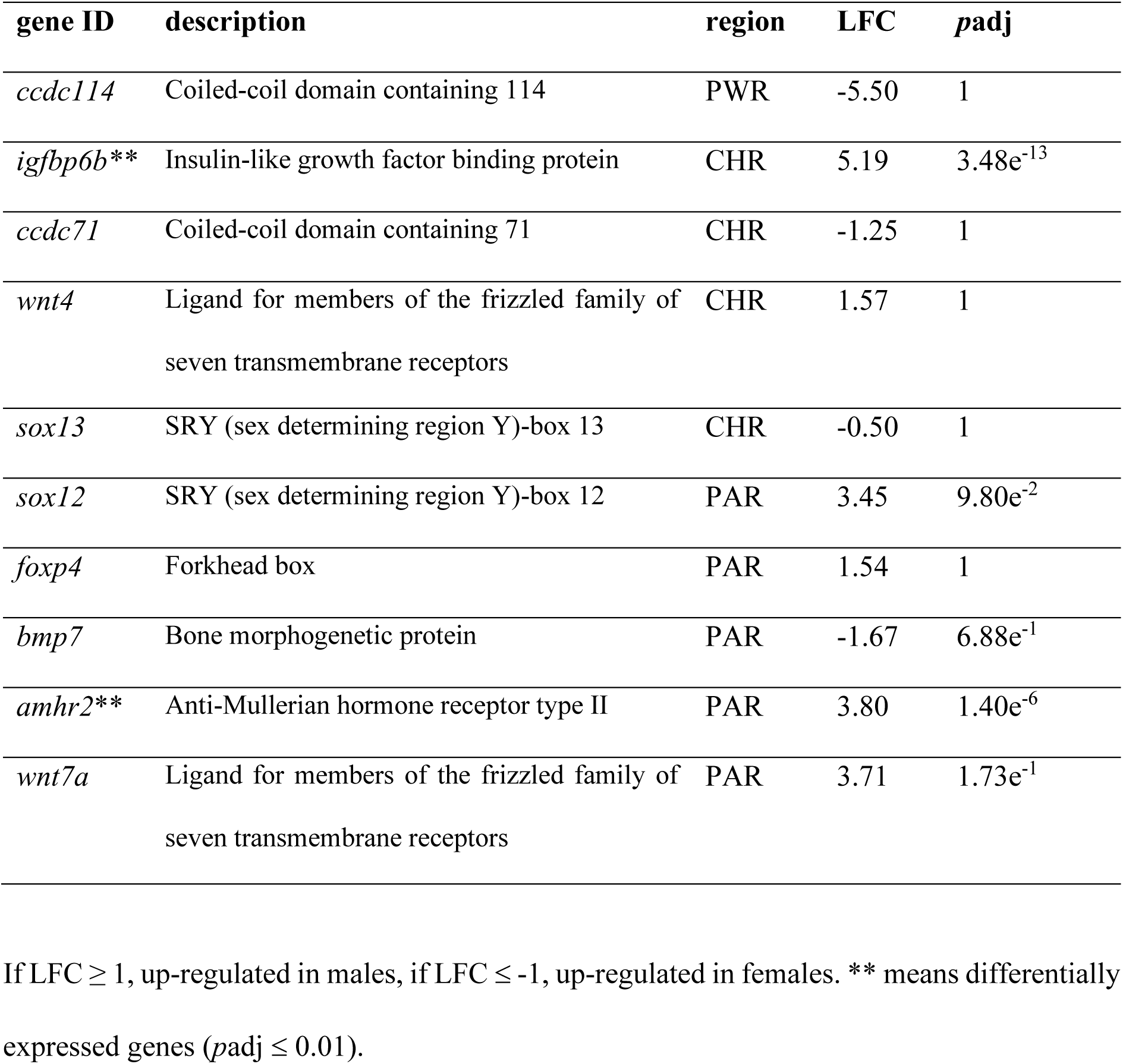
Sex determination-related genes present in *M. macrocephalus* sex chromosome.

## 3. Discussion

### 3.1 Chromosome-level genome

In this study, several strategies of genome sequencing were used to assemble a chromosome-level reference genome for *Megaleporinus macrocephalus*, a neotropical fish species with heteromorphic sex chromosomes. Despite the great morphological and ecological diversity in Characiformes [34], they have scarce genomic record.

The piauçu genome presented quality, contiguity metrics (contig and scaffold N50), and size similar to other available assemblies of Neotropical fish (**Supplementary Table 8**). The repeat content found (46.71%) in the piauçu genome was intermediate to what has been reported in other Neotropical fish species, such as *C. macropomum* (52.49%) [35] and *A. mexicanus* (41%) [36]. DNA transposons were the most abundant type of TE (11.82%), corroborating what has been observed in teleost fish [37]. A great percentage (18.89%) of the interspersed repeats remained unclassified, which was also reported in tambaqui [35] and red-bellied piranha [38] genomes (39.15% and 28.3% of unclassified sequences, respectively). In general, the repeat content found in the sex chromosome was slightly higher than in the autosomes (4.24%), which is expected. At the repeat landscape (**Supplementary Figure 1**), it is possible to observe two bursts of transposition dominated by DNA transposon, as reported in other teleost fish *e.g.*, Nile tilapia [39].

The genome annotation resulted in 30,501 protein-coding predicted genes **(Supplementary Table 3)**, which is consistent with other related neotropical fish genomes such as tambaqui [35], red-bellied piranha [40] and cavefish [41] (31,149; 30,575 and 25,293, respectively).

### 3.2 Satellite DNA

Satellite DNAs (sat DNA) are composed of arrays containing nearly identical repeating units, varying in length from single base pairs (mononucleotide repeats) to several megabases, uninterrupted [42]. The considerable length of these arrays presents a significant challenge for modern sequencing, assembly, and mapping techniques, making the analysis of lengthy fragments a formidable task [43]. The long reads are much better equipped to characterize the variable satellite content and to assemble and span difficult, repetitive parts of the genome [44]. As far as we know, piauçu has the highest number of satellites characterized for a given species so far [12]. In 2019, [12] established a satellitome library for piauçu to enhance the assembly of the sex chromosomes using long reads. However, even using this library, the amount of sat DNA found in our assembly was significantly lower than expected, which suggests that:

1. The long reads were not capable of capturing the complete arrays of these repeats. High-identity regions as the tandem repeats, often collapse during assembly with short or erroneous long reads [44] [45].
2. The pipelines used to annotate the repeats were not able to identify the satellite DNA arrays. According to [46], computational tools that take into account the high error rates of long-read technologies are lacking [46]. The employment of personalized pipelines, such as in [6] [47] [48], or tools specifically designed for satellite analysis in long-reads, like NCRF [46], tandem-genotypes [49], P ACMON STR [50], TandemTools [51] and Winnowmap2 [52], could improve the satellite DNA annotation results.
3. Sat DNAs were removed from the assembly during the scaffolding process. Conventional Hi-C analysis approaches often fail to account for reads that map to multiple locations, resulting in an underestimation of biological signals from repetitive genome regions [44] [48]. This disproportionally affects repetitive parts of the genome, such as the sex chromosomes [44] [6] [48] [53].

### 3.3 Linkage map

In the present study, we obtained a similar resolution of other linkage maps constructed for related Neotropical fish species that present analogous karyotype characteristics (haploid chromosome number, morphology, and size (**Supplementary Table 9**). The sex chromosome was syntenic with two linkage groups, LG 24 and LG 27, that respectively represent the Z-recombining region and PAR of the sex chromosome. This pattern was previously reported in the butterfly *Melitaea cinxia* [54], which also presents a ZW sex chromosome system. The LG24 was composed of markers that followed a Z chromosomal inheritance, *i.e.*, female offspring are homozygous of one of the father’s alleles. This explains the strong heterochiasmy of this LG, in which higher recombination was observed in the male map. Otherwise, although being separated by the LG24 linkage pattern, LG27 is physically merged (in the genome) to this LG and corresponds to the pseudo-autosomal region of the sex chromosome, as similarly observed in *M. cinxia* [54].

Piauçu male linkage map was longer than the female, with a genetic-length ratio of 1.07. Recently, the tambaqui *C. macropomum* was described as having an XY hypothetical sex determination system [55], despite not presenting heteromorphic sex chromosomes. [56] found that the tambaqui female linkage map was larger than the male (1.55x). Differences in map length can result from a variation in the number of recombination events in the two parents as well as variations in the number and location of the mapped loci. It is common to find a difference in the recombination ratio between the two sexes in most aquatic species [57] [58] [59] [60]. Despite this being a common phenomenon, the mechanism responsible for the different recombination rates between the sexes is still not well understood [61]. This explains the opposite sex-specific differences observed between piauçu and tambaqui and suggests that the heterogametic sex presents smaller maps due to recombination suppression [60]. The influence of the sex-determination system in the sex-specific recombination patterns was also described for other fish lineages. In flatfish, turbot [62], Senegalese sole [63], and Atlantic halibut [60], female maps were larger (1.36. 1.32. and 1.07 times); while in the Japanese flounder *Paralichthys olivaceus* [59] and tongue sole *Cynoglossus semilaevis* [61], the male maps were slightly larger (1.03 and 1.09 times, respectively).

#### 3.3.1 Inconsistencies between genetic and physical mapping

Our linkage map was successfully used as a reference to anchor the genome scaffolds into a chromosome-scale, evidencing its high quality. The chromosome-level genome anchored using the linkage map presented a high correspondence with the reference genome (scaffolded with Hi-C physical mapping). These results were similar to those obtained by other chromosome-level genomes anchored with linkage maps, such as in *A. mexicanus* [41] and *Sander lucioperca* [64]. The inconsistencies revealed by structural differences (relocations and inversions) between the linkage groups and chromosomes ordered by Hi-C (physical mapping) were also reported in the Lake Trout *Salvelinus namaycush* [65] and probably will need further investigation using other techniques, such as physical mapping of specific DNA into chromosome spreads with FISH (Fluorescence *in situ* hybridization).

### 3.4 Sex chromosome characterization

The initial long-read technologies, like PacBio CLR and Nanopore, offered more complete and contiguous genome assemblies and improved resolution of complex genomic regions. However, their high inherent error rates presented challenges in accurately distinguishing between X and Y, or Z and W haplotypes [66]. Current sex chromosome assemblies using long reads often rely on alignment with a previous reference, typically constructed with short reads, to differentiate sex chromosome-related reads/contigs, as seen in emu W [67] and threespine stickleback Y [6].

In our study, we assembled a highly degenerated sex chromosome in a non-model species without prior genomic references, utilizing PacBio long reads and Hi-C. The insufficient differentiation of piauçu Z and W chromosome sequences in the non-recombining region, coupled with the use of error-prone PacBio CLR reads, resulted in a fused sex chromosome containing a mix of Z and W sequences. A similar outcome was observed in the assembly of the threespine stickleback Y, employing the same technologies [6]. Despite the fused sex chromosome, the integration of different approaches (recombination suppression, coverage, *F_st,_* and number of SNPs) allowed the identification of 3 Mb of a W-specific region; 18 Mb of a chimeric region, constituted mostly by Z-specific sequences, but also by sequences that present allelic differences between Z and W; and 24 Mb of the pseudoautosomal region.

### 3.5 Sex chromosome gene repertoire

Genes situated in the non-recombining region, namely *ccdc114*, *igfbp6*, *sox13,* and *wnt4,* belong to gene families involved in various developmental processes in fish, including sex differentiation and determination [68]. Despite the relevance of their gene families in sex determination/differentiation, neither *ccdc114*, *igfbp6* nor *sox13* are currently recognized to actively participate in these processes or have been identified as master sex-determining (MSD) genes. While a few genes unrelated to the gonadal development process have been reported as key regulators, like *gdf6*, it is less likely that the mentioned genes are linked to piauçu sex determination. Otherwise, *wnt4* plays a crucial role in triggering the formation of ovaries during fish sex differentiation [69]. In tambaqui (*C. macropomum*), it was related to sex differentiation, either upregulated in female-like individuals or antagonized in male-like individuals [60], suggesting that it could also play a role in sex regulation and dimorphism in piauçu.

Within the recombining region, genes from the TGF-β signaling pathway were identified (*bmp7, amhr2*). Members of this signaling pathway have recurrently and independently emerged as MSD genes in vertebrates (refer to the comprehensive review in [60]). Notably, among the 20 distinct MSD genes identified so far, 13 belong to the TGF-β signaling pathway (*amh*, *amhr2*, *bmpr1b*, *gsdf*, and *gdf6*).

Bone morphogenetic proteins (BMPs) are implicated in mammalian germ cell specification and gametogenesis [70]. Recently, a truncated form of a BMP type I receptor, BMPR1BB, was identified as the MSD gene in *Atlantic herring* [7]. *bmp7* was not identified as a candidate sex-determining gene in any species so far, but it was related to sex differentiation processes in mouse embryos [71] and fish [72]; therefore, further studies should be performed to better understand the function of this gene for sex determination in piauçu.

*amhr2* is the anti-Müllerian hormone (AMH) receptor and was coopted as an MSD gene in various fish species [73] [74] [75]. Due to the relevance of the *amh*/*amhr2* pathway in sex determination, especially in fish, we highlight *amhr2* as another candidate for sex determination in piauçu. We hypothesize that a long-distance receptor located in the non-recombining region is inhibiting *amhr2* transcription, directing the sex fate toward females. A similar mechanism was reported in the Amami spine rat [76].

## 4. Methods

### 4.1 Chromosome-level genome

Tissue samples for genome sequencing were obtained from an adult ZW female of *Megaleporinus macrocephalus* from the broodstock of the Aquaculture Center of São Paulo State University. To confirm the genotype of the individual, we performed cytogenetic analysis using the lymphocyte culture technique described by [77] with some adjustments and C-banding according to Sumner, 1972 (**Supplementary Figure 5**).

To generate long reads, high molecular weight (HMW) DNA was extracted from blood using Nanobind CBB Big DNA Kit (Circulomics), and a CLR library was constructed using SMRTbell Express Template Prep Kit 2.0. The library was sequenced in one single-molecule real-time (SMRT) cell of the PacBio Sequel II System. All previous steps were performed by the Genomics & Cell Characterization Core Facility (GC3F) of the University of Oregon (USA). To improve the accuracy of the long reads, a short read library was produced with MGIEasy PCR-Free Library Prep Set (MGI Tech Co., Ltd.), sequenced on a BGI MGISEQ-2000 150 bp PE in the BGI Genomics facility. Finally, to merge the scaffolds into putative chromosomes, a chromatin interaction (Hi-C) library was generated using Proximo Hi-C Library Prep Kit (Phase Genomics) with *in vivo* cross-linking in the Genomic Sciences Laboratory of the North Carolina State University (USA). Sequencing was performed on an Illumina NovaSeq 6000 150 bp PE.

#### 4.1.1 Genome Size Estimate

The short reads were used to estimate the haploid genome size, rate of heterozygosity, and abundance of repetitive elements. First, the reads were trimmed with Trimommatic [79] [80]and bases with an average quality < 20 within a sliding window of 4 bp and bases with quality < 20 at the beginning and the end of the reads were removed. Reads with length < 36 were also discarded. After filtering, Jellyfish [81] [82] was used to count canonical *k*-mers (-C flag) of length ranging from 21 to 24. The resulting *k*-mer profile was loaded on GenomeScope [83][84].

#### 4.1.2 Genome Assembly

An initial contig assembly was performed with Falcon/Falcon-Unzip [13] [14]with a minimum read length cutoff of 5,000 bp. Falcon [13] [14] was run with default parameters, except for computing the overlaps. Raw read overlaps were computed with daligner parameters -v -k16 -w7 -h64 -e0.70 -s1000 -M27 -H5000 to better reflect the higher error rate in PacBio Sequel II. Preassembled read (pread) overlaps were computed with daligner parameters -v - k20 -w6 -h256 -e0.96 -s1000 -l2500 -M27 -H5000. Falcon-Unzip [13] [14] was run with default parameters and resulted in a set of primary and alternate contigs. False duplications in the contigs were removed with Purge_Dups [85]. Short-read polishing was made with Polca [86] [87]. To polish primary and alternate assemblies, we first concatenated them and followed with one round of short-read polishing. To improve the assembly’s contiguity, we used PacBio long reads > 10 kb to fill in spanned gaps with SAMBA [88] [87]. The Juicer [89,90] and 3d- dna pipelines [91] [90] were used to orient scaffolds into putative chromosomes. First, we generated a file with the location of DpnII enzyme restriction sites in the assembly (*generate_site_positions.py*) and a file with scaffold sizes. Second, Hi-C reads were aligned to the assembly and filtered by Juicer [92] to generate a duplicated-free list of paired alignments (merged_nodups file). At last, 3d-dna [91] was run with a minimum scaffold size of 10 kb. The resulting contact map was manually curated in Juicebox Assembly Tools (JBAT) [93] following a post-curation process. At last, we performed an extra round of polishing with Polca [86] [87].

#### 4.1.3 Quality Assessment of Genome

The correctness was evaluated in each assembly step using Merqury [23] [22]. The tool compared assembly *k*-mers to those found in the unassembled highly accurate MGISEQ short reads to estimate base-level accuracy (consensus quality value, QV) and *k*-mer completeness. The QV represents a log-scaled probability of error for the consensus base calls. Contiguity measures such as contig and scaffold N50 were obtained with the *stats.sh* script of BBMap [94]. To assess the completeness of the genome, we performed BUSCO analysis [95] [96] using the Actinopterygii dataset. The assembly was verified for contamination by the National Center for Biotechnology Information (NCBI) submission protocols. All the contaminated scaffolds identified were removed.

#### 4.1.4 Karyotype Validation

To validate the quality of our assembly, we performed a Pearson’s correlation of the estimated size in base pair (bp), based on the average karyotype size in micrometers (µm), and the assembled size (bp) of each chromosome. For this, we measured both arms of each chromosome pair of the female karyotype and calculated an average size (µm) for each chromosome. The estimated chromosome size was calculated using the formula: chromosome average size (µm) x total genome size (bp) / total karyotype size (µm).

#### 4.1.5 Repeat Annotation

We used RepeatModeler2 [97] [98], with the LTR option enabled, to produce a custom *de novo* library of the repeats present in the genome. Next, Repeat Masker [99] was used to identify, classify, and mask repetitive elements, including low-complexity sequences and interspersed repeats. We used a combined library to run Repeat Masker. First, the RepBase RepeatMasker Edition (version 20181026) was combined with the Dfam library with *addRepBase.pl* and *configure.pl* scripts. Then, only the repeats present in Teleost were selected with *famdb.py*. Finally, the custom de novo library, the Teleost repeat sequences, and a satellite library of the species [12] were concatenated. Repeat elements were soft-masked with RepeatMasker.

#### 4.1.6 Gene Prediction and Annotation

We performed gene prediction with *ab initio* and homology-based methods using the BRAKER [24] [25] pipeline. First, BRAKER1 [24] [25] used RNA-seq data (8.4.4 **Erro! Fonte de referência não encontrada.**) as extrinsic evidence to predict introns. Next, BRAKER2 [24] [25] used protein homology information from Orthodb sequences of Vertebrata. At last, TSEBRA [27] [28] selected the best transcripts from both predictions to increase their accuracies. Then, we performed a sanity check on the dataset to include only high-quality predictions. To assign functional annotation to the gene models, we performed searches using the predicted proteins with the Actinopterygii dataset of UniProtKB [29]. Search results were loaded into Blast2GO [100] [101], mapped, and annotated. The quality of the annotation was evaluated using BUSCO [102] [96].

### 4.2 Linkage mapping

To construct a linkage mapping, we produced 4 full-sib families using single mating (1 female x 1 male) during the breeding season of December 2018, totalizing 299 progeny individuals (**Supplementary Table 10**). The breeders belonged to the population kept at the Aquaculture Center of São Paulo State University (UNESP), Jaboticabal (São Paulo State, Brazil). Induced spawning was performed using carp pituitary extract dissolved in saline solution (0.9% NaCl) and applied in two dosages, with a 12 h interval: the first and second dosage of 0.6 and 5.4 mg/kg for females, and a single dosage of 1.5 mg/kg for males, at the same time of the females’ second dosage. After hatching in 20 L conical fiberglass incubators, the larvae were transferred to tanks of 250 L. The larvae were fed with *Artemia nauplii* for 20 days. Gradually, the feed was replaced by 50% of crude protein. In the fingerling stage, 1.2 mm pelleted feeds were used (40% of crude protein) and provided twice daily (commercial feed Nutripiscis Presence).

Each full-sib family was kept separately in individual fiberglass tanks of 1 m^3^ up to 6 months old. The fish were kept in a water recirculation system, fitted with mechanical and biological filters, an external aeration system, and controlled temperature at 30 °C (standard deviation = 0.5 °C) using a thermal controller connected to heaters (2 × 500 watts). Temperature, dissolved oxygen, and pH were measured with a Multiparameter Water Quality Checker U-50 (Horiba). After this period, we collected blood samples for genomic analyses, and the weight of all animals was registered with analytical balance (average weight was 6 g). Fish were then euthanized for sex identification. Individual sex was verified by a PCR-based protocol using a chromosome W-probe [12] as well as by cytogenetic analysis. Chromosome preparations were obtained from kidney tissues using the technique described by [103].

#### 4.2.1 SNP genotyping

DNA was extracted from blood samples with Wizard Genomic DNA Purification kit (Promega) and quality was verified in 1% agarose gel electrophoresis. Purity was accessed in Nanodrop One and concentration (ng/μl) was measured by Qubit fluorometer with Qubit dsDNA HS Assay kit (Invitrogen, USA). We used a modified version of the protocol described by [104] for the construction of ddRADseq libraries. Briefly, 75 ng of genomic DNA from each individual was digested (8 U/reaction) using the combination of two restriction enzymes, SphI and MluCI (New England Biolabs), and ligated to specific adapters (P1 and P2, 0.25 μM) using the enzyme T4 DNA ligase, at 23°C for one and a half hour and 65°C for 10 minutes to heat kill the enzyme. The P1 adapters have an additional 5 nucleotides that function as individual tags (barcode). The selection of digested fragments was performed using E-Gel Power Snap System (Thermo Fisher Scientific) with fragments of approximately 350 bp. Subsequently, PCR assays were performed to incorporate the identification of each library. In total, 7 libraries were constructed, with an average of 46 samples/library. PCR was performed under the conditions of the Platinum SuperFi DNA Polymerase enzyme (Thermo Fischer Scientific). The reactions were purified with the ProNex Size-Selective Purification System kit (Promega) and the concentration was rechecked by fluorometry in the Qubit 3.0 instrument (Thermo Fisher Scientific). Finally, the libraries were sequenced in 2 lanes of Illumina Hiseq2500 150 PE, using 15 % PhiX (Novogene).

The overall quality of raw sequencing data was checked using FastQC [105]. Next, the data were analyzed using Stacks [106] [107] for SNPs calling. Briefly, sequences were demultiplexed and filtered using *process_radtags* and individual reads that passed the previous quality filters were aligned to the chromosome-level reference genome of *M. macrocephalus*. Subsequently, *gstacks* created loci by incorporating the ddRAD-aligned reads. Finally, *populations* was used to generate genotype data for the samples. To differentiate putative SNPs from sequencing errors, we used Plink 1.9 [30] [31] to filter spurious SNPs with more than 10% genotyping error rate (--geno 0.1), minor allele frequencies less than 0.05 (--min-maf 0.05), and Hardy-Weinberg imbalance (*p* < 5E10-5). Regarding the removal of individuals, samples that had more than 15% (--mind 0.15) of absent genotypes were excluded.

#### 4.2.2 Linkage map

A linkage map was created using Lep-MAP3 [108] [109]. First, a parenthood test was performed using the *IBD* module, and individuals with more than 10% of Mendelian errors were removed. The *ParentCall2* module was used to impute possible missing genotypes or to correct erroneous parental genotypes based on progeny data. *Filtering2* module was used to remove markers with significant segregation distortion (dataTolerance = 0.001) and non-informative markers. Markers were assigned to LG by *SeparateChromosomes2* using the minimum LOD score. The best LOD was selected iteratively and accounted for marker distribution in the first 27 linkage groups, which corresponds to the haploid chromosome number of the species. Next, orphan markers were assigned to existing linkage groups (LOD score lower than in *SeparateChromosomes2*) using *JoinSingles2* and ordered within each linkage group using the *OrderMarkers2* module. Due to the slight stochastic variation in marker distances between runs, the *OrderMarkers2* module was run 15 times and the order with the best likelihood value for each LG was selected.

The reliability of the SNP *loci* attribution on the LG and the respective *loci* ordering within the LGs was verified through comparative genomic synteny analysis with the reference genome using *Circa* [110].

We used the genome scaffolds to generate other chromosome-level genome using the linkage map as a reference in Chromonomer [32] [33]. This was done to verify possible differences between the linkage map ordering (genetic mapping) and the Hi-C ordering (physical mapping). Chromonomer [32] [33] attempts to find the best set of nonconflicting markers that maximizes the number of scaffolds in the resulting genome while minimizing ordering discrepancies. It resulted in a FASTA file (chromonome.fa), the chromosome-level genome oriented according to the genetic map.

### 4.3 Resequencing (pool-sequencing)

We used resequencing analyses to contrast whole-genome sex differences in *M. macrocephalus*. For this purpose, we collected samples of 20 males and 20 females originating from four commercial fish farms in Brazil. Briefly, fish were anesthetized with 0.1% benzocaine for blood collection. The sex of each fish was verified by cytogenetic analysis, as detailed above in **5.2 Linkage mapping**, and samples were clustered in separated male and female pools.

DNA was extracted individually and quantified according to the **5.2 Linkage mapping** section and next clustered in male and female pools. Library construction and sequencing were performed at INRAE (Rennes, France) in the Laboratory of Physiology and Genomics of Fish (LPGP) using an Illumina NovaSeq S4 platform 150 bp PE.

The Pool-Seq dataset was analyzed with the Pooled Sequencing Analysis for Sex Signal (PSASS) pipeline [111]. Briefly, reads from the male and female pools were mapped into the female pseudo-haplotype chromosome-level genome (GCA_021613375.1) using bwa-mem [112] [113] with default parameters. Then, the alignment files were sorted, merged and PCR duplicates were removed with Picard tools [114]. Reads with mapping quality < 20 and that were not mapped uniquely were also removed with samtools [115] [116]. Next, the two sex BAM files were used to generate a pileup file using samtools mpileup [115] [116] with per-base alignment quality disabled (−B). A sync file was created using popoolation mpileup2sync (parameters: --min-qual 20) [117], which contained the nucleotide composition of each sex for each position in the reference genome. With this sync file, *F*_ST_, SNPs, and coverage between the two sexes in all reference positions were calculated in a 50 kb sliding window with an output point every 1,000 bp to identify sex-specific SNPs enriched regions.

### 4.4 RNA-seq

For RNA-seq experiments, 60 individuals that comprised the offspring of one full-sib family of *M. macrocephalus* were used. Fish were produced and maintained as described above (**5.2 Linkage mapping**). At 150 days after fertilization, when the period of sex differentiation recently occurred according to previous experiments in this species *(unpublished data*); the two gonads and kidneys of each fish were dissected immediately. Fish were euthanized by benzocaine anesthetic overdose (2%) for sampling. One gonad was stored in RNAlater (Thermo Fischer Scientific) for RNA extraction, and the other was fixed for 24 hours in Karnovsky’s solution [118] and then stored in ethanol 70% for phenotypic sex identification in microscopy. The sex of each fish was verified by cytogenetic analysis, as detailed above in **5.2 Linkage mapping.** The phenotypic sex was obtained through gonadal histology as described by [119].

After phenotypic and genotypic sex identification, the samples were clustered in two pools: ZZ males and ZW females. Each pool had three biological replicates that consisted of 10 gonads, resulting in 6 libraries for RNA sequencing. RNA was extracted from each pool with RNeasy Micro Kit (Qiagen). Next, the integrity (RIN > 7) and concentration (ng/µl) were accessed using Bioanalyzer 2100 (Agilent). At last, library construction and sequencing were performed by BGI Genomics (Shenzhen, China) using the BGISEQ-500 platform 100 bp PE.

Raw read quality was accessed using FastQC [105]. Adapters and poor-quality reads were trimmed in Trimmomatic [79] [80] (parameters LEADING:20 TRAILING:20 SLIDINGWINDOW:4:20 MINLEN:36). Trimmed reads were pseudo-aligned against mRNA sequences obtained from *M. macrocephalus* genome (GCA_021613375.1) with kallisto [120] [121]. A matrix with estimated counts of transcripts abundance was exported with R/tximport [122] [123]. Differential expression analysis was performed with R/DESeq2 [124] [125], using the design formula ∼ sex. Transcripts with False Discovery Rate (FDR) adjusted *p*-values ≤ 0.05 were considered differentially expressed. Transcripts with Log Fold Change (LFC) ≥ 1 were considered as up-regulated in males and transcripts with LFC ≤ −1 were up-regulated in females.

## Abbreviations

BMPs: Bone morphogenetic proteins
BUSCO: Benchmarking Universal Single Copy Orthologs
CLR: continuous long reads
ddRADseq: Double Digest Restriction Site Associated DNA Sequencing
DE: differentially expressed
FISH: Fluorescence in situ hybridization
*F_st_*: fixation index
GO: gene ontology
HMW: high molecular weight
JBAT: Juicebox Assembly Tools
LG: Linkage Group
LINE: long interspersed nuclear elements
LOD: Logarithm of Odds
LTR: long terminal repeats
MSD: master sex determining
NCBI: National Center for Biotechnology Information
PacBio: Pacific Biosciences
PAR: Pseudoautosomal region
PCA: Principal component analysis
QV: Consensus Quality Value
sat DNA: Satellite DNAs
SINE: short interspersed nuclear elements
SMRT: single molecule real time
T2T: Telomere to Telomere consortium
TE: transposable elements.

## Additional Files

**supplementary_material_tables.docx** – Supplementary tables.

**supplementary_material_figures.pdf** – Supplementary figures.

**supplementary_material_gene_description.xlsx** – Table containing gene ID, chromosome, position (bp), gene symbol and description.

**supplementary_material_DE_results.xlsx** – Table containing Differential Expression analysis results (gene ID, log_2_ fold change, *p*adj, chromosome, position (bp), gene symbol and description).

## Declarations

### Ethics approval

This study was conducted in strict accordance with the recommendations of the National Council for Control of Animal Experimentation (CONCEA) (Brazilian Ministry of Science, Technology, and Innovation) and was approved by the Ethics Committee on Animal Use (CEUA number 4936/20) of Faculdade de Ciências Agrárias e Veterinárias, UNESP, Campus Jaboticabal, SP, Brazil.

### Availability of data and materials

This Whole Genome Shotgun project has been deposited at DDBJ/ENA/GenBank under the accession JAJQXZ000000000. The version described in this paper is version JAJQXZ010000000. The assembled genome is available at the NCBI with the accession number GCA_021613375.1.

### Authors’ contributions

DTH, RU and CHSB conceived and designed the study. DTH, RU, RH, YG, FPF, FF, AB and CP supervised the research. CHSB wrote the manuscript with inputs from DTH. CHSB, MUS, AV and DTH performed bioinformatic analysis. DTH, RU, FPF, FF, RH and YG provided funding. CHSB constructed the ddRADseq libraries. CHSB and SM extracted DNA of pool-sex samples. CHSB, DTH, JFGA, LVGL, MVF, RBA performed induced spawning of breeders for the ddRADseq experiment. CHSB, DTH, RU and RBA performed cytogenetics analysis. CHSB, AJB and LVGL performed histologic analysis and RNA extraction of RNAseq samples. RSH, AJB, CHSB and LVGL analyzed the histology samples. CHSB, DTH, RU, JFGA, MVF, VAMF, RBA collected data and samples. All authors read and approved the final manuscript.

## Acknowledgments

This research article was possible thanks to the scholarship granted from the Brazilian Federal Agency for Support and Evaluation of Graduate Education (CAPES), in the scope of the Program CAPES-PrInt, process number 88887.467255/2019-00. Also, we would like to thank Valdecir Fernandes de Lima and Marcio Roberto Reche for their support in collecting the samples and handling the fish.

