## Supplementary Material for "De novo assembly and characterization of a highly degenerated ZW sex chromosome in the fish Megaleporinus macrocephalus"

**Supplementary Table 1.** Assembled and estimated chromosomes sizes (bp) calculated using karyotype data.

| Autosome | Assembled size (bp) | Estimated size (bp) |
|---|---|---|
| 1 | 73,843,892 | 73,524,246 |
| 2 | 57,073,771 | 58,279,251 |
| 3 | 54,318,282 | 52,872,252 |
| 4 | 53,925,970 | 52,480,849 |
| 5 | 53,011,304 | 51,891,237 |
| 6 | 52,262,800 | 51,390,179 |
| 7 | 51,642,936 | 49,819,898 |
| 8 | 49,518,186 | 49,300,333 |
| 9 | 49,298,696 | 48,760,706 |
| 10 | 48,705,945 | 48,068,703 |
| 11 | 48,550,980 | 48,064,379 |
| 12 | 47,357,970 | 47,656,718 |
| 14 | 44,903,196 | 47,083,191 |
| 15 | 44,847,474 | 47,013,143 |
| 16 | 44,428,828 | 46,953,300 |
| 17 | 44,393,724 | 46,065,509 |
| 18 | 44,386,724 | 44,600,904 |
| 19 | 43,994,550 | 43,546,728 |
| 20 | 43,858,395 | 42,646,484 |
| 21 | 43,700,133 | 41,890,833 |
| 22 | 41,427,154 | 41,198,311 |
| 23 | 40,492,453 | 41,016,705 |
| 24 | 40,120,662 | 40,253,271 |
| 25 | 40,052,844 | 40,168,695 |
| 26 | 38,727,815 | 39,137,695 |
| 27 | 36,538,851 | 38,981,687 |
| **Pearson** | **0,99** | |

**Supplementary Table 2.** Repeat annotation statistics for *Megaleporinus macrocephalus* genome.

| | **No. of elements** | **Length (bp)** | **% of genome sequence** |
|---|---|---|---|
| **Retroelements** | 325,354 | 86,900,511 | 6.78 |
| SINEs: | 32,263 | 4,249,898 | 0.33 |
| Penelope | 1,992 | 200,769 | 0.02 |
| LINEs: | 135,677 | 43,882,452 | 3.42 |
| L2/CR1/Rex | 109,233 | 36,159,292 | 2.82 |
| R1/LOA/Jockey | 538 | 129,887 | 0.01 |
| R2/R4/NeSL | 222 | 69,099 | 0.01 |
| RTE/Bov-B | 10,276 | 2,603,408 | 0.20 |
| L1/CIN4 | 10,838 | 3,907,815 | 0.30 |
| LTR elements: | 157,414 | 38,768,161 | 3.02 |
| BEL/Pao | 2,206 | 845,394 | 0.07 |
| Ty1/Copia | 421 | 198,340 | 0.02 |
| Gypsy/DIRS1 | 24,957 | 7,950,002 | 0.62 |
| Retroviral | 10,995 | 2,687,534 | 0.21 |
| **DNA transposons** | 817,030 | 151,501,452 | 11.82 |
| hobo-Activator | 295,340 | 56,805,942 | 4.43 |
| Tc1-IS630-Pogo | 253,381 | 56,461,503 | 4.40 |
| PiggyBac | 3,214 | 595,853 | 0.05 |
| Tourist/Harbinger | 34,271 | 6,807,497 | 0.53 |
| Other (Mirage, P-element, Transib) | 333 | 20,280 | 0.00 |
| Rolling-circles | 18,420 | 4248,326 | 0.33 |
| **Unclassified:** | 1379224 | 242,219,098 | 18.89 |
| **Total interspersed repeats:** | | 480,621,061 | 37.49 |
| **Small RNA:** | 2375 | 321,707 | 0.03 |
| **Satellites:** | 88059 | 56,424,759 | 4.40 |
| **Simple repeats:** | 580511 | 52,069,796 | 4.06 |
| **Low complexity:** | 55387 | 5,150,938 | 0.40 |

**Supplementary Table 3.** Summary of the annotated features of *Megaleporinus macrocephalus* genome.

| Feature | *Megaleporinus macrocephalus* |
|---|---|
| **Genes and Pseudogenes** | 30,501 |
| **Protein-Coding** | 30,501 |
| **Exon** | 248,235 |
| **Intron** | 217,739 |
| **CDS** | 248,235 |
| **mRNA** | 30,501 |
| **Start codon** | 30,487 |
| **Stop codon** | 30, 488 |
| **Mean intron per gene** | 7.14 |
| **Mean exon per gene** | 8.14 |

**Supplementary Table 4**. Summary of ddRAD sequencing statistics.

| Library | Total sequences | Filtered sequences (%) | | | | Retained reads (%) | Average retained reads/ ind. (million) |
|---|---|---|---|---|---|---|---|
| | | Adapter sequence | No barcode | Low quality | No rad cutside | | |
| **1** | 198,679,472 | 1.19 | 3.67 | 2.37 | 1.33 | 91.44 | 3.95 |
| **2** | 191,095,010 | 1.18 | 4.92 | 2.43 | 1.66 | 89.82 | 3.73 |
| **3** | 183,389,588 | 1.15 | 4.36 | 2.59 | 1.86 | 90.04 | 3.59 |
| **4** | 188,028,608 | 1.17 | 4.38 | 2.53 | 1.81 | 90.12 | 3.68 |
| **5** | 165,196,674 | 1.09 | 4.53 | 2.42 | 1.70 | 90.27 | 3.24 |
| **6** | 207,410,674 | 1.15 | 6.49 | 2.32 | 1.86 | 88.18 | 3.98 |
| **7** | 173,700,306 | 1.15 | 6.88 | 2.76 | 2.07 | 87.14 | 3.29 |
| **Total** | 1,307,500,332 | 1.15 | 5.03 | 2.48 | 1.75 | 89.58 | 25.46 |

**Supplementary Table 5**. Summary of the genetic map of piauçu. Chr represents the chromosome which the linkage group had synteny with. Size is related to the length in bp, after scaffolding with Chromonomer. N is the number of markers. Length in cM, Density in cM/Locus.

| LG | Chr | Size | n | Sex-averaged | | Male | | Female | | M:F |
|---|---|---|---|---|---|---|---|---|---|---|
| | | | | Length | Density | Length | Density | Length | Density | |
| **1** | 6 | 23,599,497 | 710 | 126.42 | 0.18 | 135.77 | 0.19 | 131.68 | 0.19 | 1.03 |
| **2** | 4 | 45,953,104 | 688 | 130.95 | 0.19 | 127.07 | 0.18 | 107.22 | 0.16 | 1.19 |
| **3** | 1 | 49,978,048 | 625 | 133.56 | 0.21 | 139.93 | 0.22 | 130.11 | 0.21 | 1.08 |
| **4** | 15 | 40,070,618 | 543 | 120.91 | 0.22 | 139.69 | 0.26 | 133.3 | 0.25 | 1.05 |
| **5** | 10 | 43,303,417 | 476 | 122.43 | 0.26 | 106.81 | 0.22 | 100.3 | 0.21 | 1.06 |
| **6** | 18 | 34,104,434 | 484 | 112.29 | 0.23 | 135.78 | 0.28 | 115.72 | 0.24 | 1.17 |
| **7** | 2 | 50,146,178 | 489 | 129.75 | 0.27 | 156.05 | 0.32 | 121.16 | 0.25 | 1.29 |
| **8** | 9 | 44,463,537 | 503 | 107.79 | 0.21 | 116.85 | 0.23 | 115.32 | 0.23 | 1.01 |
| **9** | 25 | 36,288,099 | 429 | 135.49 | 0.32 | 137.46 | 0.32 | 135.06 | 0.31 | 1.02 |
| **10** | 23 | 36,576,158 | 501 | 122.64 | 0.24 | 152.38 | 0.3 | 117.92 | 0.24 | 1.29 |
| **11** | 16 | 32,187,476 | 444 | 111.72 | 0.25 | 108.1 | 0.24 | 80.47 | 0.18 | 1.34 |
| **12** | 8 | 43,378,694 | 442 | 127.77 | 0.29 | 136.22 | 0.31 | 99.65 | 0.23 | 1.37 |
| **13** | 11 | 36,977,433 | 473 | 121.11 | 0.26 | 128.1 | 0.27 | 124.15 | 0.26 | 1.03 |
| **14** | 12 | 44,454,369 | 406 | 141.52 | 0.35 | 142.35 | 0.35 | 135.23 | 0.33 | 1.05 |
| **15** | 27 | 35,428,719 | 381 | 119.88 | 0.31 | 120.71 | 0.32 | 119.61 | 0.31 | 1.01 |
| **16** | 17 | 38,667,919 | 372 | 139.89 | 0.38 | 139.95 | 0.38 | 141.94 | 0.38 | 0.99 |
| **17** | 19 | 25,433,675 | 385 | 121.22 | 0.31 | 124.13 | 0.32 | 133.74 | 0.35 | 0.93 |
| **18** | 24 | 19,470,523 | 339 | 119.57 | 0.35 | 121.55 | 0.36 | 121.27 | 0.36 | 1 |
| **19** | 22 | 27,071,565 | 351 | 119.33 | 0.34 | 137.5 | 0.39 | 121.55 | 0.35 | 1.13 |
| **20** | 3 | 38,970,815 | 296 | 82.15 | 0.28 | 94.36 | 0.32 | 95.58 | 0.32 | 0.99 |
| **21** | 7 | 18,150,482 | 253 | 81.57 | 0.32 | 96.9 | 0.38 | 92.98 | 0.37 | 1.04 |
| **22** | 20 | 39,059,067 | 236 | 143.08 | 0.61 | 142.45 | 0.6 | 130.32 | 0.55 | 1.09 |
| **23** | 21 | 26,787,905 | 250 | 107.64 | 0.43 | 111.77 | 0.45 | 104.11 | 0.42 | 1.07 |
| **24** | 13 | 4,059,682 | 225 | 43.25 | 0.19 | 47.37 | 0.21 | 87.81 | 0.39 | 0.54 |

| | | | | | | | | | | |
|---|---|---|---|---|---|---|---|---|---|---|
| **25** | 5 | 23,599,497 | 237 | 132.10 | 0.56 | 150.95 | 0.64 | 139.78 | 0.59 | 1.08 |
| **26** | 14 | 45,953,104 | 239 | 136.79 | 0.57 | 135.83 | 0.57 | 147.43 | 0.62 | 0.92 |
| **27** | 13 | 22,889,721 | 251 | 120.66 | 0.48 | 117.89 | 0.47 | 113.62 | 0.45 | 1.04 |
| **28** | 26 | 29,510,817 | 203 | 108.88 | 0.54 | 114.34 | 0.56 | 104.98 | 0.52 | 1.09 |
| **Total** | | 977,082,963 | 11,231 | 3320.36 | 0.29 | 3,518.24 | 0.31 | 3,301.97 | 0.29 | 1.07 |

**Supplementary Table 6**. Summary of RNA-sequencing statistics.

| Pool | Replicate | Total reads | Retained reads (%) | Total data (Gb) |
|---|---|---|---|---|
| | 1 | 46,331,866 | 97.72 | 4.53 |
| **Male** | 2 | 50,141,272 | 97.93 | 4.91 |
| | 3 | 41,915,706 | 100.00 | 4.44 |
| | 1 | 50,886,670 | 97.58 | 4.97 |
| **Female** | 2 | 48,466,868 | 97.21 | 4.71 |
| | 3 | 48,337,546 | 97.33 | 4.70 |
| | **Total** | 286,079,928 | 97.91 | 28.26 |

**Supplementary Table 7**. Top 10 genes up-regulated in females ZW (pink) and males ZZ (blue).

| transcript ID | LFC | *p*adj | chr | gene ID | description |
|---|---|---|---|---|---|
| Mmac_g35407.t1 | -12.73 | 2.57E-13 | chr6 | - | Zona pellucida sperm-binding protein 3-like |
| Mmac_g39735.t2 | -12.28 | 2.79E-82 | chr9 | - | Zona pellucida sperm-binding protein 3-like |
| Mmac_g34977.t1 | -11.75 | 1.19E-08 | chr6 | *Zp4* | Zona pellucida |
| Mmac_g34976.t2 | -11.57 | 3.05E-14 | chr6 | *Zp4* | Zona pellucida |
| Mmac_g21232.t2 | -11.54 | 1.54E-05 | chr21 | - | Zona pellucida sperm-binding protein 3-like |
| Mmac_g36656.t1 | -11.24 | 1.26E-61 | chr7 | - | Zona pellucida sperm-binding protein 3-like |
| Mmac_g18381.t1 | -11.13 | 2.41E-33 | chr2 | - | Zona pellucida sperm-binding protein 3-like |
| Mmac_g14296.t1 | -10.99 | 2.45E-58 | chr18 | - | Zona pellucida sperm-binding protein 3-like |
| Mmac_g36128.t1 | -10.92 | 1.07E-16 | chr7 | *Aqp1* | Belongs to the MIP aquaporin (TC 1.A.8) family |
| Mmac_g7248.t1 | -10.88 | 3.09E-12 | chr13 | *Smarcd1* | SWI SNF related, matrix associated, actin dependent regulator of chromatin, subfamily d, member 1 |
| Mmac_g2999.t1 | 9.31 | 1.75E-54 | chr10 | *Ces5a* | Belongs to the type-B carboxylesterase lipase family |
| Mmac_g33548.t1 | 6.67 | 3.80E-32 | chr5 | *Enpep* | Glutamyl aminopeptidase |
| Mmac_g22869.t1 | 8.50 | 6.87E-26 | chr22 | - | Insulin / insulin-like growth factor / relaxin family. |
| Mmac_g38128.t1 | 9.00 | 4.30E-24 | chr8 | *Ppp1r1c* | Protein phosphatase 1 regulatory |
| Mmac_g5032.t1 | 6.79 | 1.53E-23 | chr11 | *Coch* | Coagulation factor C homolog. cochlin (*Limulus polyphemus*) |
| Mmac_g12538.t1 | 6.78 | 1.53E-23 | chr16 | - | Gonadal somatic cell derived factor |
| Mmac_g4256.t1 | 7.03 | 1.55E-22 | chr11 | *Fabp7* | Belongs to the calycin superfamily. Fatty-acid binding protein (FABP) family |
| Mmac_g19985.t1 | 6.30 | 7.59E-21 | chr20 | *Amh* | Anti-mullerian hormone |
| Mmac_g39628.t1 | 6.39 | 3.34E-20 | chr9 | - | Apelin receptor |
| Mmac_g25145.t1 | 6.20 | 1.14E-18 | chr24 | *Tbx1* | T-box transcription factor |

**Supplementary Table 8**. Summary of sampling number per family and sex

| Family | Males | Females | Total |
|---|---|---|---|
| **1** | 49 | 44 | 93 |
| **2** | 4 | 20 | 24 |
| **3** | 49 | 44 | 93 |
| **4** | 41 | 48 | 89 |
| **Total** | 143 | 156 | 299 |

**Supplementary Table 9.** Comparison between available genome assemblies of Neotropical fish species

| **Species** | **Genome size*(bp)** | **Scaffold N50 (bp)** | **Contig N50 (bp)** | **Haploid chromosome number (n)** |
|---|---|---|---|---|
| *Megaleporinus macrocephalus* (GCA_021613375.1) | 1,280,781,66 | 45,034,219 | 5,013,076 | 27 |
| *Colossoma macropomum* (GCA_904425465.1) | 1,221,809,066 | 40,163,545 | 5,645,235 | - |
| *Pygocentrus nattereri* (GCA_015220715.1) | 1,222,050,449 | 42,283,192 | 12,898,870 | 30 |
| *Astyanax mexicanus* (GCA_000372685.2) | 1,291,596,431 | 35,377,769 | 1,767,240 | 25 |

*Ungapped length

**Supplementary Table 10.** Statistics of a few Neotropical fish species linkage maps.

| Species | Technique | No. of SNPs | Length (cM) | Average marker interval (cM) | Reference |
|---|---|---|---|---|---|
| *Megaleporinus macrocephalus* | ddRADseq | 11,231 | 3,320.36 | 0.29 | This study |
| *Colossoma macropomum* | GBS* | 7,734 | 2,811 | 0.39 | Nunes *et al.*, 2017 |
| | RADseq | 14,805 | 2,752 | 0.51 | Varela et al., 2021 |
| *Piaractus mesopotamicus* | RADseq | 17,453 | 2,755.60 | 0.47 | Mastrochirico-Filho *et al.*, 2020 |

*Genotype-by-sequencing

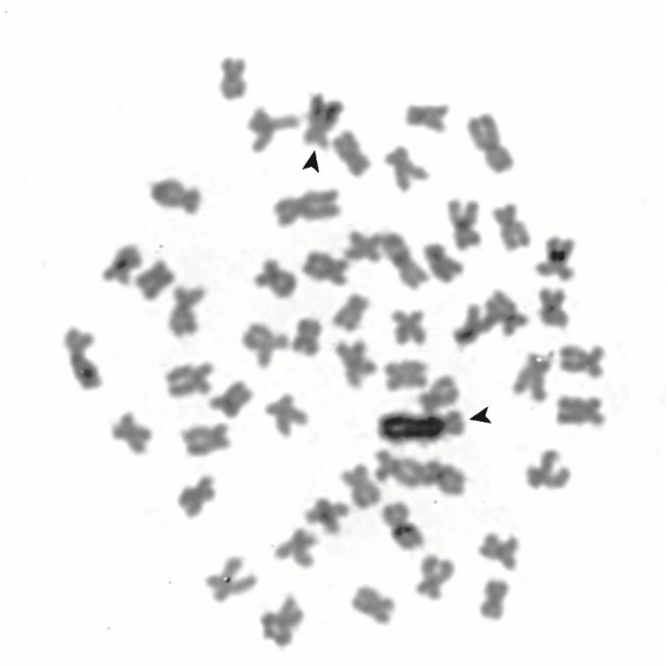

**Supplementary Figure 1**. Karyotype of a female of *Megaleporinus macrocephalus* under C-banding. Arrows are indicating Z and W sex chromosomes.

.

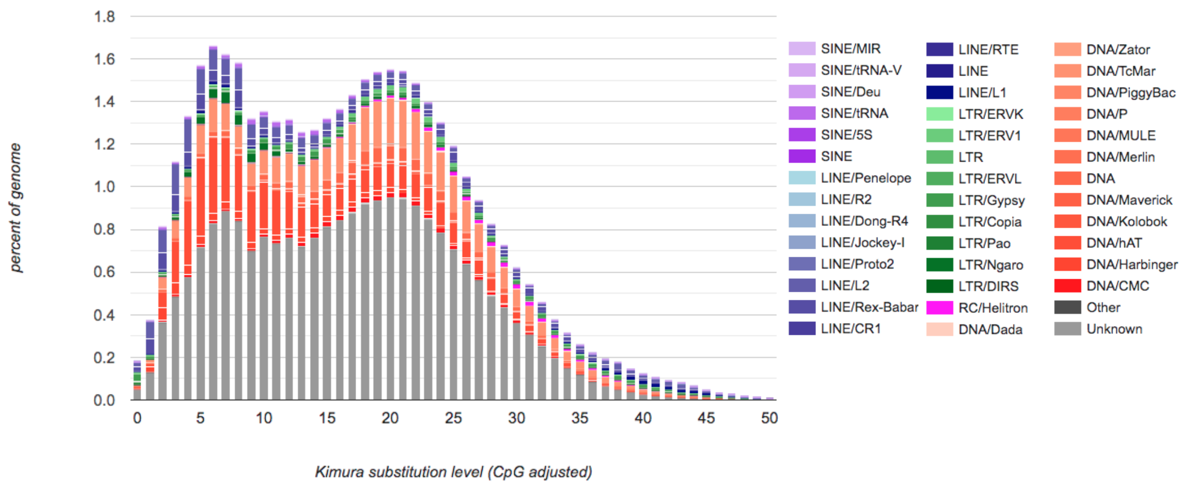


**Supplementary Figure 2.** Interspersed repeat landscape of *Megaleporinus macrocephalus* genome. The graph represents genome coverage (*y* axis) for each type of TEs (DNA transposons, SINE, LINE, and LTR retrotransposons), clustered according to Kimura distances (*K*-value; Kimura, 1980) to their corresponding consensus sequence (*x* axis, *K*-values from 0 to 50).

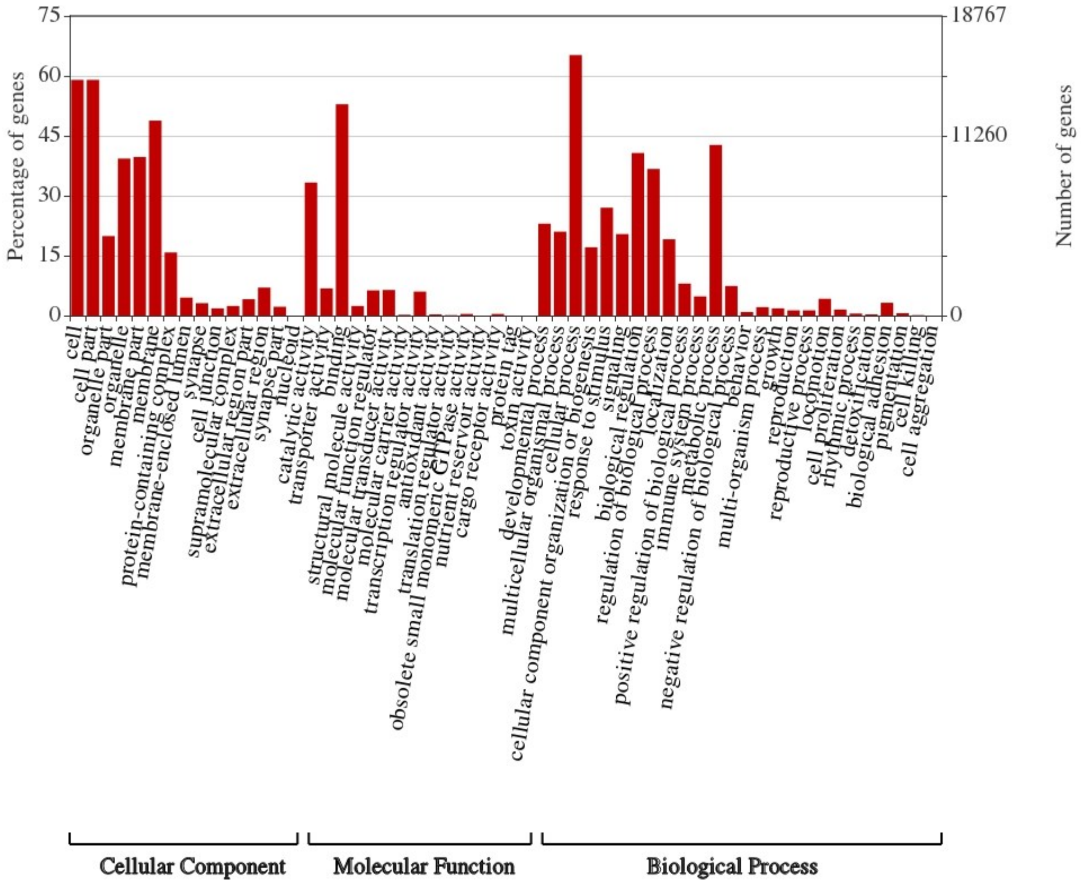


**Supplementary Figure 3.** Gene Ontology (G.O) Terms of Cellular Component, Molecular Function and Biological Process domains of *Megaleporinus macrocephalus* genome.

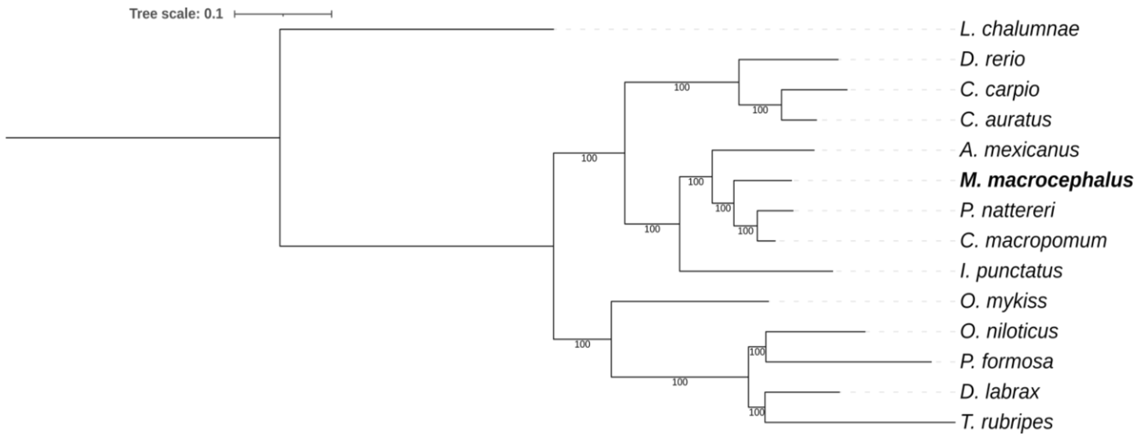


**Supplementary Figure 4**. Phylogeny of single-copy orthologs genes (n=336) of *Megaleporinus macrocephalus* and other fish species using the criteria of Maximum Likelihood. Bootstrap values are shown below the corresponding branches.

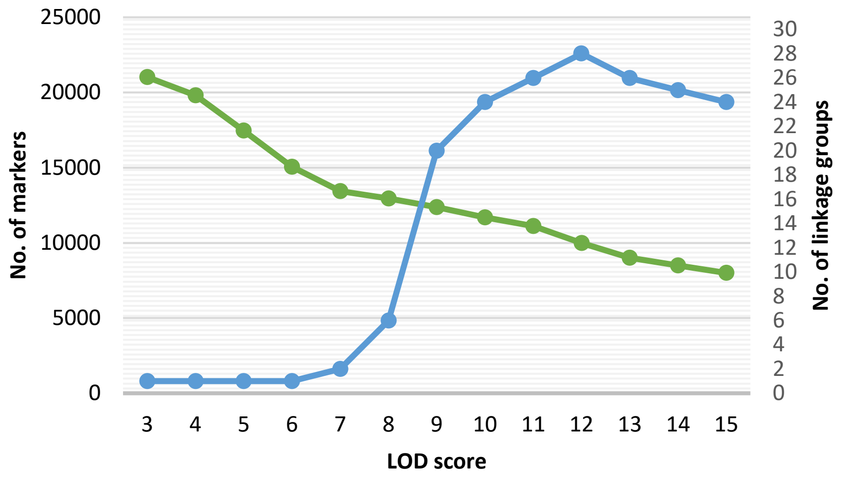


**Supplementary Figure 5.** The average number of markers in linkage groups (left y axis) and the number of linkage groups (right *y* axis) according to LOD score (*x* axis).

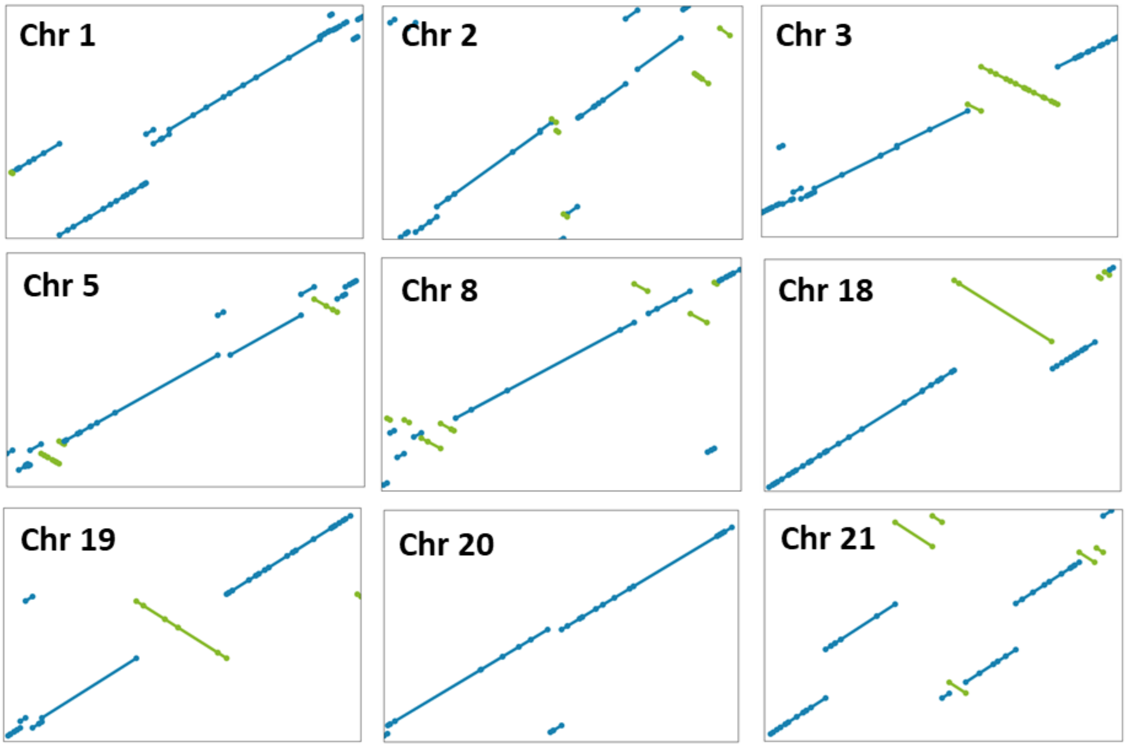


**Supplementary Figure 6.** Dotplot synteny between chromosomes constructed with Hi-C data (*x*-axis) and scaffolds of the linkage groups (*y*-axis). In blue, forward alignments; and in green, reverse alignments (inversions). The dots represent the end of scaffolds.
